# Targeting C5aR1 Increases the Therapeutic Window of Radiotherapy

**DOI:** 10.1101/2020.10.27.358036

**Authors:** Monica M. Olcina, Melemenidis Stavros, Dhanya K. Nambiar, Ryan K. Kim, Kerriann M. Casey, von Eyben Rie, Trent M. Woodruff, Edward G. Graves, Le Quynh-Thu, Stucki Manuel, Amato J. Giaccia

## Abstract

Engaging innate immune pathways is emerging as a productive way of achieving durable anti-tumor responses. However, systemic administration of these therapies can result in toxicity, deemed to be particularly problematic when combined with current standard-of-care cytotoxic treatments such as radiotherapy. Increasing the therapeutic window of radiotherapy may be achieved by using targeted therapies, however, few pre-clinical studies investigate both tumor and normal tissue responses in detail. Here we show that targeting innate immune receptor C5aR1 improves tumor radiation response while reducing radiation-induced normal tissue toxicity, thereby increasing the therapeutic window. Genetically or pharmacologically targeting C5aR1 increases both IL-10 expression in the small intestine and IL-10 secretion by tumor cells. Increased IL-10 attenuates RelA phosphorylation and increases apoptosis in tumor cells, leading to improved radiation responses in murine models. Of note, these radiosensitizing effects are tumor-specific since, in the gastrointestinal tract, targeting C5aR1 instead results in decreased crypt cell apoptosis reduced signs of histological damage and improved survival following total abdominal irradiation in mice. Furthermore, the potent and orally active C5aR1 inhibitor, PMX205, improves tumor radiation responses even in a context of reduced/absent CD8+ T cell infiltration. These data indicate that PMX205 can modulate cancer-cell intrinsic functions to potentiate anti-tumor radiation responses even in tumors displaying features of T-cell deficiency or exclusion. Finally, using a preclinical murine model allowing the simultaneous assessment of tumor and normal tissue radiation responses, we show that PMX205 treatment reduces histological and functional markers of small-bowel toxicity while affording a positive tumor response. Our data, therefore, suggest that targeting C5aR1 could be a promising approach for increasing the therapeutic window of radiotherapy.

## Introduction

As a first responder to danger, the innate immune system senses pathogen- or danger-associated molecular patterns and initiates an inflammatory response to clear the danger (Burdette et al., 2011). Cytotoxic therapies such as radiotherapy can induce danger-associated molecular patterns, and thus stimulating sensor innate immunity pathways together with radiotherapy has been investigated as a means of improving anti-tumor responses (Dar et al., 2019; Ricklin et al., 2016). This approach has led to some impressive pre-clinical improved radiation responses, for example using STING or TLR agonists in combination with radiotherapy (Dar et al., 2019; Dewan et al., 2012; Liang et al., 2017). However, many of these approaches still rely on significant anti-tumor immune infiltration/ T-cell responses for their full effectiveness (Baird et al., 2016; Dewan et al., 2012; Liang et al., 2017). Furthermore, systemic administration of these therapies unfortunately often leads to significant side effects (Su et al., 2019). An unfavorable side effect profile might be particularly problematic in the context of radiotherapy, which itself leads to radiation-induced normal tissue toxicity (Hauer-Jensen et al., 2014; Leibowitz et al., 2014). For example, pelvic irradiation will be delivered for up to 300,000 patients worldwide with 50-70% of these estimated to experience acute toxicity (Andreyev, 2007; Ashburn and Kalady, 2016; Zimmerer et al., 2008). The small intestine is often subject to some of the most distressing damage when incidentally present in the field during pelvic, upper gastrointestinal tract, inferior lung or retroperitoneal irradiation (Ashburn and Kalady, 2016; Kavanagh et al., 2010). The resulting condition is known as radiation-induced small bowel disease (also historically known as radiation enteritis or enteropathy) (Ashburn and Kalady, 2016). In fact, in clinical practice, minimizing toxicity associated with radiotherapy while still achieving maximal tumor control is actively considered when preparing treatment plans (Baumann et al., 2016; Bentzen, 2006; Giaccia, 2014). Despite the importance of considering both tumor and normal tissue responses in clinical practice, few preclinical studies have comprehensively investigated the effects of radiation-modulating therapies on both tumor and normal tissue responses (Moding et al., 2013). Given the important role of the immune system in maintaining normal tissue homeostasis, it seems important to consider immune targets that can enhance tumor radiation responses without negatively impacting normal tissues in the irradiation field.

The complement system is a key innate immunity pathway that serves to sense, recognize and clear foreign pathogens and altered self (Gros et al., 2008; Pio et al., 2013; Ricklin et al., 2010). Targeting the complement system is emerging as a promising therapeutic approach in cancer, however, whether targeting the pathway together with radiotherapy is beneficial or detrimental remains unclear (Elvington et al., 2014; Reis et al., 2017; Surace et al., 2015). Furthermore, the effects of targeting this pathway on radiation-induced normal tissue toxicity have not been evaluated. Proteolytic cleavage of complement components leads to their activation and the formation of downstream effectors (Sayegh et al., 2014). Complement cleavage products signal through cell surface receptors such as C3aR1 and C5aR1. C5aR1 is the main receptor for C5a, which is considered the most potent anaphylatoxin of the complement cascade (Sayegh et al., 2014). C5aR1 is the most advanced complement target in the oncology space with preclinical and clinical trial efforts mainly focused on blocking C5aR1 to improve anti-tumor immunity (Ajona et al., 2017; Markiewski et al., 2008; Massard et al., 2019; Wang et al., 2016). However, given the recently discovered autocrine and intracellular cancer-cell intrinsic roles of complement system proteins, it is important to also investigate the contribution of these functions to tumor responses (Bai et al., 2019; Block et al., 2019; Cho et al., 2014; Olcina et al., 2020). The discovery of cancer cell intrinsic functions that can potentiate anti-tumor responses without having to rely on increased T-cell infiltration would be particularly beneficial, especially in the context of so-called “cold” tumors (Duan et al., 2020; Roumenina et al., 2019).

Here we report that targeting C5aR1 with a potent noncompetitive orally active inhibitor, PMX205, results in improved tumor radiation responses following increased IL-10-dependent apoptosis. Interestingly, we find that these effects can occur *in vitro* and in tumors displaying reduced/absent CD8+ T cell infiltration, suggesting that PMX205 can modulate cancer-cell intrinsic functions that potentiate anti-tumor radiation responses without having to rely on increased T-cell numbers. Of note, these effects are tumor-specific since, in the gastrointestinal tract, targeting C5aR1 instead results in decreased apoptosis and improved survival of mice following total abdominal irradiation. These data suggest that increased apoptosis following modulation of the C5aR1-IL-10 axis is tumor-specific and can thus be advantageously targeted for therapeutic intent. Indeed, when assessing the utility of PMX205 in a combined preclincial model of tumor and normal tissue toxicity, we find that PMX205 treated mice have reduced histological and functional signs of normal tissue toxicity and the best tumor responses. Taken together, these results suggest that C5aR1 is a good target for improving the therapeutic index of radiotherapy by increasing tumor radiosensitivity and decreasing normal intestinal tissue toxicity.

## Results

### C5aR1 as a good therapeutic target

Targeting the complement system is emerging as a popular strategy to enhance anti-tumor immunity (Ajona et al., 2017; Elvington et al., 2014; Markiewski et al., 2008; Wang et al., 2016). However, while this is no doubt a powerful approach in terms of improving anti-tumor responses, little is known about which members should be targeted in order to improve tumor response while also minimizing normal tissue toxicity. We hypothesized that the best therapeutic candidates might be genes that affect cell survival in a stress as well as cancer-specific manner and that are amenable to pharmacological intervention. To search for these genes, we performed an *in silico* screen by mining the DepMap database, established to identify cancer dependencies by combining data from CRISPR-Cas9 and RNAi screens in more than 700 cell lines. To identify stress- and cancer-specific dependencies, we applied stringent criteria and looked for genes that were deemed non-essential across all cell lines (genes that when knocked-out/down under unstressed conditions do not alter cell survival) while still being expressed in most tissues. We reasoned that looking for non-essential hits would allow the identification of genes providing stress-specific cancer dependencies and, therefore, potential therapeutic targets less likely to mediate toxicity in normal tissues upon depletion. Furthermore, a gene was only considered a hit if it was “druggable” based on structural and ligand-based assessment (as assessed in canSAR). Genes that are hits and are druggable are shown in green. When querying complement system components, receptors, proteases and regulators as reported in (Ricklin et al., 2010) we found three hits: *C5*, *C5AR1*, *C4BPA*. *ATR* was included in the screen a positive control for essential genes since its deletion is lethal is several cell lines due to its role in replication and the DNA damage response (Cimprich and Cortez, 2008; Flynn and Zou, 2011). Interestingly, *C1QBP,* a complement gene which was recently shown to play a role in the DNA damage response by modulating DNA resection, was essential in a number of cell lines, further validating our screening approach (Bai et al., 2019)(**Figure 1A**) **(Supplementary Table 1)**.

**Figure 1.**
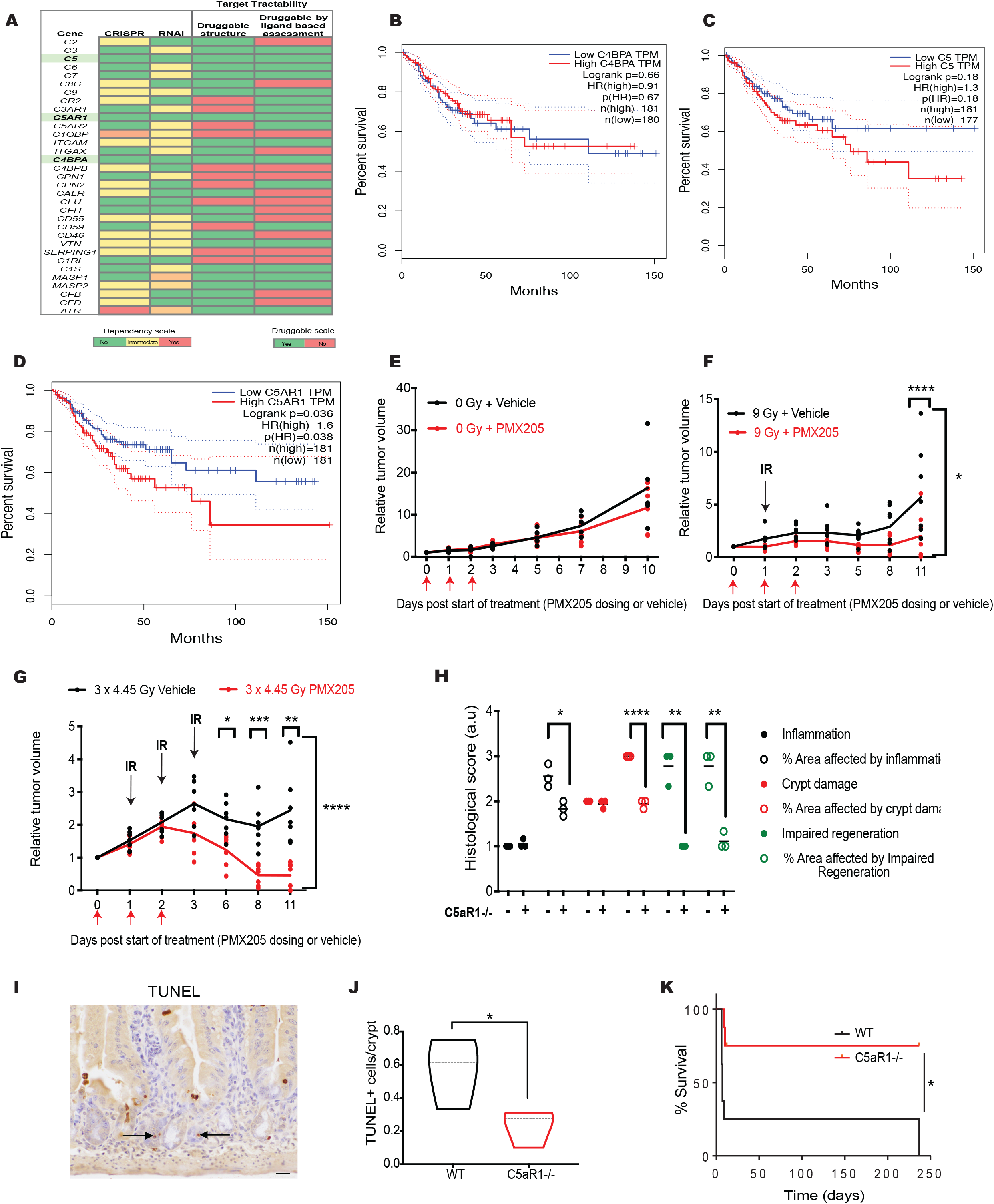
C5aR1 as a good therapeutic target. (A) *In silico* screen of complement genes. Data acquired from DepMap portal (https://depmap.org) Essential genes are shown in red. Non-essential genes across all studies are shown in green. Yellow/Orange = intermediate dependence or essentiality. For target tractability: green corresponds to druggable structure = Yes and druggable by ligand-based assessment = Yes. Red corresponds to druggable structure = No and druggable by ligand-based assessment = No. (B) GEPIA Kaplan-Meier (KM) curve for disease-free survival of TCGA colorectal cancer patients with high (red) or low (blue) C4BPA mRNA expression levels is shown. Group Cutoff = Median (http://gepia.cancer-pku.cn). (C) GEPIA Kaplan-Meier (KM) curve for disease-free survival of TCGA colorectal cancer patients with high (red) or low (blue) C5 mRNA expression levels is shown. Group Cutoff = Median (http://gepia.cancer-pku.cn). (D) GEPIA Kaplan-Meier (KM) curve for disease-free survival of TCGA colorectal cancer patients with high (red) or low (blue) C5AR1 mRNA expression levels is shown. Group Cutoff = Median (http://gepia.cancer-pku.cn). (E) Relative tumor growth curves are shown for MC38 subcutaneous tumors treated with either vehicle or PMX205 treatment for 3 doses (on day 0, 1 and 2). (F) Relative tumor growth curves are shown for MC38 subcutaneous tumors treated with 9 Gy single dose irradiation and either vehicle or PMX205 treatment for 3 doses flanking the irradiation dose (on day 0, 1 and 2). **** = p<0.0001 comparing vehicle and PMX205 treated mice at day 11. * = p<0.05 comparing day 0 to day 11 by 2-way ANOVA with Dunnett’s comparison test. Individual points represent individual mice per group. (G) Relative tumor growth curves are shown for MC38 subcutaneous tumors treated with 3 x 4.45 Gy single dose (equivalent to 9 Gy assuming an α/β ratio of 5.06)(Suwinski et al., 2007) irradiation (IR) and either vehicle or PMX205 treatment for 3 doses flanking the irradiation dose (on day 0, 1 and 2). * = p<0.05, ** = p<0.01, *** = p<0.001, comparing vehicle and PMX205 treated mice at days 6, 8 and 11 respectively. **** = p<0.0001 comparing day 0 to day 11 by 2-way ANOVA with Dunnett’s comparison test. Individual points represent individual mice per group. (H) Graph shows the breakdown of individual scores used to calculate total histological damage scores for WT and C5aR1^−/−^ mice irradiated with 9 Gy total abdominal irradiation. Intestines were harvested 2 days post-IR. Slides scored blindly by a pathologist. **** = p<0.0001, ** = p<0.01 2-tailed t-test. Individual points represent individual mice per group. (I) Representative image of TUNEL immunoreactivity (arrows) in a WT section is shown. (J) Graph shows the number of TUNEL + cells in WT or C5aR1^−/−^ mice irradiated with 9 Gy total abdominal irradiation. Intestines were harvested 2 days post-IR. ** = p<0.01, 2-tailed t-test. Individual points represent individual mice per group. (K) Kaplan-Meier curve for WT or C5aR1-/- mice irradiated with 12 Gy total abdominal irradiation. * = p<0.05, 2-tailed t-test, n = 7 per group.

*C5* encodes complement component C5, which when cleaved following complement cascade activation will form C5a, the most potent anaphylatoxin of the complement system. C5aR1 is the main receptor for the C5a ligand. Targeting the C5a/C5aR1 axis was recently shown to enhance anti-tumor immune responses, fueling interest in testing this approach in clinical trials (Ajona et al., 2017; Markiewski et al., 2008; Massard et al., 2019; Wang et al., 2016). *C4BPA* encodes the alpha chain of complement regulator C4BP (Hofmeyer et al., 2013). We recently found that *C4BPA* mutations are associated with improved overall survival across epithelial cancers and certain *C4BPA* mutations confer sensitivity to standard-of-care chemotherapy agents used in colorectal cancer treatment (Olcina et al., 2020). However, no known pharmacological approaches for targeting C4BPA have been reported. To further narrow down which hit would be the best therapeutic target, we assessed the association of C5, C5aR1 and C4BPA mRNA expression with prognostic outcomes and found that only high C5aR1 mRNA expression was associated with significantly poor disease-free survival in colorectal cancer (and overall survival in all TCGA cancer types queried) **(Figure 1B-D and Supplementary Figures 1A and B)**. We, therefore, decided to pursue the investigation of C5aR1 as a target for enhancing the therapeutic window of radiotherapy and to investigate its effects on both tumor and normal tissue. To do so, we first investigated the impact of C5aR1 depletion on tumor response with the use of C5aR1^−/−^ mice and a syngeneic colorectal cancer model. Remarkably, following intraperitoneal (i.p.) injection of CT26 colorectal cancer cells, there was reduced tumor burden in the C5aR1^−/−^ mice compared to wild-type (WT) controls, although these differences did not reach statistical significance likely due to the high variability in tumor weights across mice. Irradiation of WT or C5aR1^−/−^ mice injected with CT26 cells also reduced tumor burden in both cases but only the C5aR1^−/−^ mice had undetectable tumor burden in this model and also, consequently, statistically significantly lower tumor weight compared to WT mice **(Supplementary Figures 1C-E).** While these data highlight the importance of host C5aR1 on tumor uptake, they also suggest that this is not a good model for specifically assessing the effect of targeting C5aR1 on tumor radiation response. We, therefore, decided to move into alternative syngeneic models and to use pharmacological rather than genetic inhibition of C5aR1 in subcutaneous tumor models. This approach would allow us to treat tumors of equivalent initial tumor volume and to directly interrogate the effect of targeting C5aR1 either in the absence of irradiation or in combination with different irradiation fractionation regimens while following tumor growth over time. For these studies we decided to use PMX205, a cyclic hexapeptide potent noncompetitive inhibitor of C5aR1 suitable for oral dosing (Kumar et al., 2020). PMX205 has been previously used in preclinical models and found to be a specific C5aR1 antagonist (Jain et al., 2013; Kumar et al., 2018, 2020; Lee et al., 2017; Li et al., 2020). Furthermore, PMX205 has been granted Food and Drug Administration (FDA) and European Medicines Agency (EMA) “orphan drug” status allowing accelerated progression to clinical trials (Ricklin and Lambris, 2016). In MC38 subcutaneous tumors, treatment with three doses of C5aR1 antagonist PMX205 did not have significant effects on tumor response in the absence of irradiation and had no negative effects on mouse weight **(Figure 1E and Supplementary Figure 1F)**. We then decided to investigate the effects of single dose (9 Gy) and equivalent multiple fractionation (3 × 4.45 Gy) regimens (equivalent assuming an α/β ratio of 5.06) (Suwinski et al., 2007) together with PMX205 treatment flanking the irradiation doses. We found that both fractionation and single dose irradiation were more effective following treatment with PMX205 (following short PMX205 dosing just flanking the irradiation doses) **(Figures 1F and G)**.

We then assessed the effect of targeting C5aR1 in radiation-induced normal tissue toxicity. To this end, we established a histologic scoring system, whereby increasing doses of radiation-induced small intestinal damage were assessed through scoring of 6 different histologic parameters **(Supplementary Table 2).** Briefly, histologic parameters included neutrophilic inflammation (and distribution), crypt damage (and distribution), and crypt regeneration impairment (and distribution). All slides were blindly evaluated by a board-certified veterinary pathologist. We confirmed the validity of our damage scoring system by irradiating mice with increasing doses of total abdominal irradiation. As shown in **Supplementary Figure 1G,** increasing doses of irradiation resulted in corresponding increases in total damage scores within the small intestine, validating our scoring system. We then used the scoring system to assess whether differences in radiation-induced histologic damage could be observed between WT and C5aR1^−/−^ mice. Interestingly, we observed that C5aR1^−/−^ displayed a significantly decreased total damage score relative to WT mice **(Supplementary Figure 1H)**. The difference in the score was evident even when all six parameters were assessed separately with the greatest differences being observed in % crypt damage area and impaired regeneration scores **(Figure 1H)**. Of note, the total crypt damage score was the same between the two groups of mice, consistent with the fact that both groups received the same irradiation dose **(Figure 1H)**. Also, the same scores were observed along the length of the small intestine, with equivalent crypt damage, for instance, observed in the duodenum, jejunum and ileum **(Supplementary Figures 1I-K)**. Intrigued by the striking reduction in radiation-induced small bowel toxicity (as assessed by histology) in the C5aR1^−/−^ compared to WT mice, we assessed other markers of radiation-induced small bowel toxicity, such as vessel density, apoptosis and Ki67 staining **(Figures 1I, J and Supplementary Figures L-O)**. Interestingly we only found changes in apoptosis between WT and C5aR1^−/−^ **(Figures 1I and J).** Furthermore, the differences in apoptosis were observed primarily in the crypts and not villi consistent with the crypt harboring the most radiosensitive cell population **(Supplementary Figure O)**. Finally, we compared survival in WT and C5aR1^−/−^ mice following a higher irradiation dose of 12 Gy (expected to kill >90% of WT mice) and found that survival was significantly improved in C5aR1^−/−^ compared to WT mice **(Figure 1K).** These data indicate that C5aR1 depletion in mice results in decreased small intestinal histologic damage score, decreased intestinal crypt cell apoptosis and increased survival following total abdominal irradiation.

Together these data suggest that C5aR1 is a good therapeutic target that may increase the therapeutic window of radiation.

### Targeting C5aR1 can mediate an improved radiation response in the context of reduced CD8+ T-cell infiltration

Our results showing that absence of host C5aR1 results in reduced tumor implantation and tumor burden following irradiation **(Supplementary Figures 1C-E)** suggest that C5aR1 depletion might alter host immunity to promote anti-tumor immune responses. This would be in line with previous reports highlighting C5aR1’s role in mediating an immunosuppressive tumor microenvironment and in modulating anti-tumor CD8+ T-cell responses (Ajona et al., 2017; Markiewski et al., 2008; Wang et al., 2016). We, therefore, decided to investigate whether tumor immune infiltration changes following PMX205 and irradiation could explain the improved tumor radiation response observed. Interestingly, although 9 Gy irradiation, as expected, significantly increased levels of CD3+ and CD8+ T-cells; treatment with PMX205 did not further increase the % of these cells in the tumor **(Figure 2A, B and Supplementary Figure 2A)**. In fact, the % of CD8+ T-cells was significantly reduced in PMX205 treated mice receiving irradiation compared to vehicle treated mice **(Figure 2B)**. No significant changes in NK cells or either monocytic or granulocytic myeloid derived suppressor (M-MDSC or PMN-MDSC respectively) cells were observed between treatment groups **(Figure 2C-F)**. We also investigated whether there were any changes in immune populations in the spleens of these tumor-bearing mice and noted that all treatment groups had comparable levels of all immune populations investigated **(Figure 2G and H and Supplementary Figures 2A-F)**. The lack of tumor immune infiltration changes that could account for the improved radiation response observed in our subcutaneous tumor models was puzzling, and suggested that additional, potentially T-cell independent mechanisms could be mediating the improved response. In agreement with this hypothesis, moderate tumor growth delays were also observed in athymic mice harboring HCT 116 xenografts treated with PMX205 and 9 Gy **(Supplementary Figure 2G)**. In order to investigate which additional immune-independent mechanisms were contributing to the improved radiation response we decided to investigate whether modulation of pro-survival signaling pathways might be altered following C5aR1 inhibition. We first assessed AKT signaling since C5a/C5aR1 signaling was previously shown to modulate AKT phosphorylation (Cho et al., 2014). We observed a slight decrease in AKT phosphorylation at serine 473 (AKT-S473) following PMX205 treatment and irradiation **(Supplementary Figure 2H)**. We also asked whether PMX205 might affect p53 activity but did not observe any changes in p53 phosphorylation at serine 15 (p53-S15) **(Supplementary Figure 2H)**. Finally, we assessed whether targeting C5aR1 might affect NF-κB signaling since we recently reported that complement protein C4BPA-NF-κB crosstalk regulates apoptosis in colorectal cancer cells (Olcina et al., 2020). RelA phosphorylation at serine 536 (RelA-S536) increases transcriptional activation of RelA (Jiang et al., 2003; Madrid et al., 2001; Zhong et al., 2002). Interestingly, we observed a robust decrease in RelA-S536 following PMX205 treatment **(Figure 2I)**. Since we had observed significantly decreased apoptosis following genetic C5aR1 targeting in normal tissue **(Figure 1J)** and NF-κB is a key modulator of apoptosis following radiotherapy, we decided to assess apoptosis levels in cancer cells (Chen et al., 2000; Wang et al., 2004). Interestingly, and in contrast to what we had observed in intestinal crypts, genetically and pharmacologically targeting C5aR1 resulted in significantly increased apoptosis in colorectal cancer cells **(Figure 2J, K and Supplementary Figure 2I).** Increased apoptosis following PMX205 treatment is in line with the improved radiation responses observed in PMX205 treated tumors and indeed we observed increased apoptosis in HCT 116 xenografts treated with PMX205 and 9 Gy **(Figure 2L)**. Intrigued by the opposing effects on apoptosis observed following C5aR1 targeting in tumor and normal tissues we assessed whether C5aR1 might also alter NF-κB signaling in normal tissues, or whether changes in NF-κB signaling are cancer-cell specific. In the intestinal epithelium, ionizing radiation has been shown to activate NF-κB signaling within 0.5-1 hours following irradiation which is thought to serve as a protective mechanism from irradiation-induced apoptosis (Egan et al., 2004; Wang et al., 2004). We therefore asked whether there were any changes in RelA phosphorylation 1 hour post irradiation, when RelA phosphorylation should be near maximal levels *in vivo* (Egan et al., 2004). Here, we observed a trend towards increased RelA phosphorylation between WT and C5aR1^−/−^ mice consistent with a decrease in apoptosis in the crypts (**Supplementary Figures 2J-L).** We also investigated whether there were any differences in CD45+ or CD8+ T cells between WT and C5aR1^−/−^ mice and noted that while total CD45+ cell numbers were comparable between the two groups of mice, CD8+ T cells were reduced in the intestines of C5aR1^−/−^ mice, similarly to what had been observed in the tumors treated with PMX205 (**Supplementary Figures 2M and N).** These data suggest that increased apoptosis following modulation of the C5aR1-NF-κB axis is tumor-specific and can thus be advantageously targeted for therapeutic intent even in a context of reduced CD8+ T-cell infiltration.

**Figure 2.**
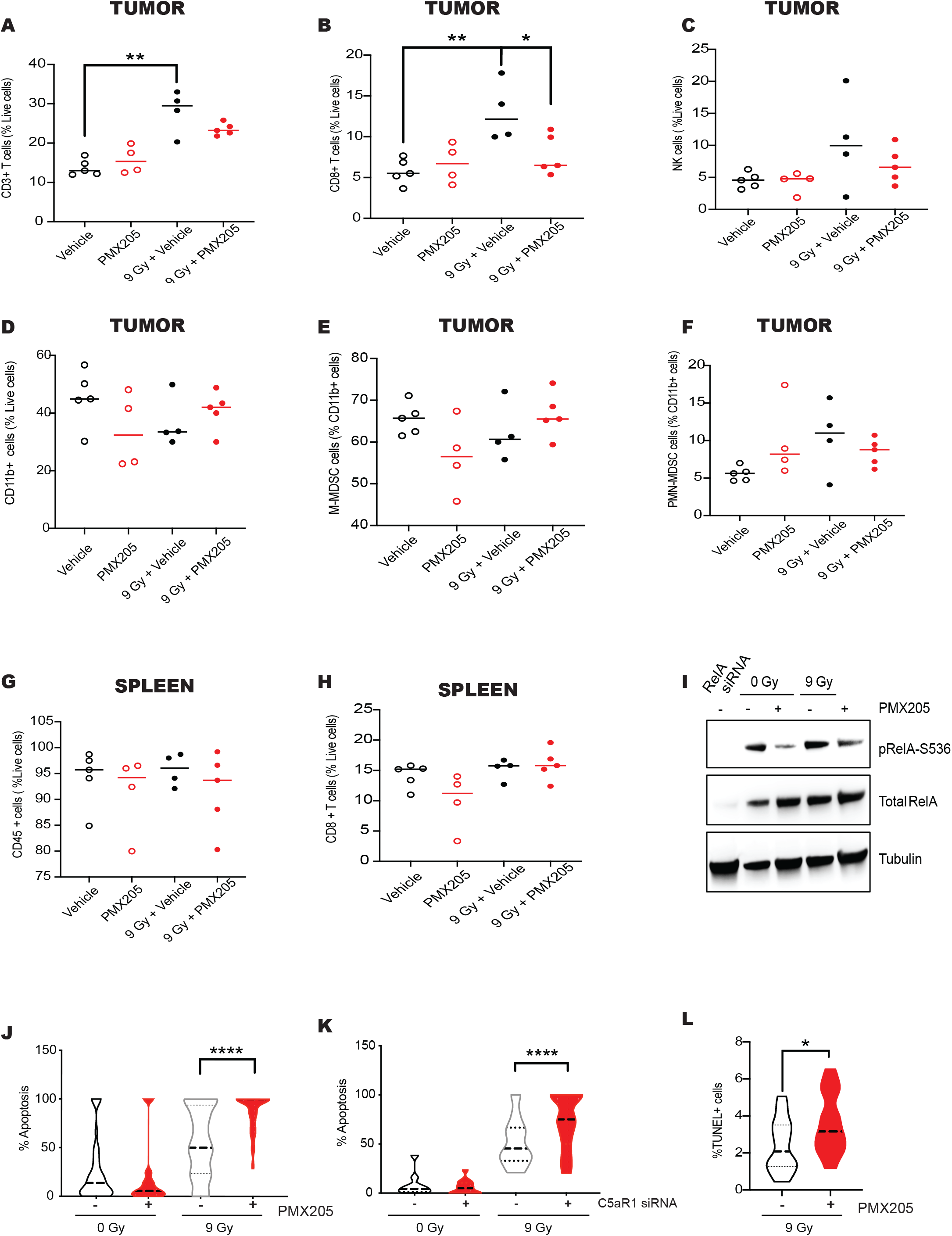
Targeting C5aR1 can mediate an improved radiation response in the context of reduced CD8^+^ T-cell infiltration. (A) Graph shows CD3+ T-cells (as a % of live cells) in tumors of mice receiving 0 or 9 Gy and either vehicle or PMX205 treatment following the same dosing scheme as shown in Figure 1F. Tumors were harvested 7 days after irradiation with either 0 or 9 Gy. ** = p<0.01 comparing 0 Gy vehicle and 9 Gy vehicle,2-tailed t-test. Individual points represent individual mice per group. (B) Graph shows CD8+ T-cells (as a % of live cells) in tumors of mice receiving 0 or 9 Gy and either vehicle or PMX205 treatment following the same dosing scheme as shown in Figure 1F. Tumors were harvested 7 days after irradiation with either 0 or 9 Gy. ** = p<0.01 comparing 0 Gy vehicle and 9 Gy vehicle, * = p<0.05 comparing 9 Gy vehicle and 9 Gy PMX205, 2-tailed t-test. Individual points represent individual mice per group. (C) Graph shows NK cells (as a % of live cells) in tumors of mice receiving 0 or 9 Gy and either vehicle or PMX205 treatment following the same dosing scheme as shown in Figure 1F. Tumors were harvested 7 days after irradiation with either 0 or 9 Gy. Individual points represent individual mice per group. (D) Graph shows CD11b+ cells (as a % of live cells) in tumors of mice receiving 0 or 9 Gy and either vehicle or PMX205 treatment following the same dosing scheme as shown in Figure 1F. Tumors were harvested 7 days after irradiation with either 0 or 9 Gy. Individual points represent individual mice per group. (E) Graph shows M-MDSC-cells (as a % of live cells) in tumors of mice receiving 0 or 9 Gy and either vehicle or PMX205 treatment following the same dosing scheme as shown in Figure 1F. Tumors were harvested 7 days after irradiation with either 0 or 9 Gy. Individual points represent individual mice per group. (F) Graph shows PMN-MDSC cells (as a % of live cells) in tumors of mice receiving 0 or 9 Gy and either vehicle or PMX205 treatment following the same dosing scheme as shown in Figure 1F. Tumors were harvested 7 days after irradiation with either 0 or 9 Gy. Individual points represent individual mice per group. (G) Graph shows CD45+ cells (as a % of live cells) in the spleen of mice receiving 0 or 9 Gy and either vehicle or PMX205 treatment following the same dosing scheme as shown in Figure 1F. Tumors were harvested 7 days after irradiation with either 0 or 9 Gy. Individual points represent individual mice per group. (H) Graph shows CD8+ T-cells (as a % of live cells) in the spleen of mice receiving 0 or 9 Gy and either vehicle or PMX205 treatment following the same dosing scheme as shown in Figure 1F. Tumors were harvested 7 days after irradiation with either 0 or 9 Gy. Individual points represent individual mice per group. (I) HCT 116 cells were treated with 0 or 9 Gy and either vehicle or PMX205 from 1 hour before irradiation (IR). Cells were harvested 48 hours post-IR. Western blotting was carried with the antibodies indicated. Tubulin was used as the loading control. n=3. Lysates from cells where RelA had been depleted by siRNA are shown to the left as a control for the detection of RelA and pRelA. (J) The graph represents the number of apoptotic/non-apoptotic cells expressed as a % of the whole population for HCT 116 cells treated with either vehicle or PMX205 from 1 hour before irradiation (IR) with either 0 or 9 Gy. Cells were harvested 48 hours post-IR. n=3. **** = p<0.0001 by 2-way ANOVA with Dunnett’s comparison test. (K) The graph represents the number of apoptotic/non-apoptotic cells expressed as a % of the whole population for HCT 116 cells treated with either Scr or C5aR1 siRNA and either 0 or 9 Gy. Cells were harvested 48 hours post-IR. n=2. **** = p<0.0001 by 2-way ANOVA with Dunnett’s comparison test. (L) The graph represents the number of TUNEL positive (+) cells found in HCT 116 xenografts (grown in athymic nude mice from the same experiment as in Supplementary Figure 2G) treated with either vehicle or PMX205 and 9 Gy, on the day of and 1 day post-irradiation. Individual points represent independent fields of view from up to 3 different tumors per group.

### Targeting C5aR1 increases apoptosis and mediates improved tumor response in an IL-10-dependent manner

Reduced NF-κB signaling following cytotoxic stress in cells harboring dysregulated complement proteins is associated with increased expression of NF-κB inhibitor, IκBα (Olcina et al., 2020). Treatment of colorectal cancer cells with PMX205, however, did not result in robust changes in IκBα expression. We also did not observe any changes in the expression of the catalytic subunit of the IKK complex, IKKα **(Supplementary Figures 3A).** These data suggest that altered NF-κB signaling downstream of PMX205 treatment occurs via a different mechanism to that previously observed in cells harboring certain complement mutations (Olcina et al., 2020).

Interestingly, autocrine complement was recently shown to inhibit production of immunosuppressive chemokine IL-10 and IL-10 is also a known radiosensitizer since it results in reduced NF-κB signaling in colorectal cancer cells (Voboril and Weberova-Voborilova, 2007; Wang et al., 2016). Furthermore, IL-10 signaling in CX3CR1+ macrophages is important for intestinal injury defense (Shouval et al., 2014; Zigmond et al., 2014). We hypothesized that targeting C5aR1 might result in increased IL-10 expression, which could be contributing to the improved tumor and normal tissue radiation responses. In order to test this hypothesis, we first assessed levels of IL-10 secretion in C5aR1^−/−^ mice and observed increased IL-10 baseline expression in the small intestines of C5aR1^−/−^ compared to WT mice when assessed by two different ELISA detection methods **(Supplementary Figure 3B)**. In contrast, following irradiation, we found that IL-10 levels were comparable in the small intestines of C5aR1^−/−^ and WT mice **(Supplementary Figure 3C)**. In the intestine, IL-10 can be produced by both epithelial as well as immune (CD45+) cells such as macrophages, which are important mediators of gut homeostasis (Shouval et al., 2014; Zigmond et al., 2014). We indeed found that both CD45- and CD45+ cells in the small intestines of WT and C5aR1^−/−^ mice expressed IL-10 **(Figure 3A and Supplementary Figure 3D)**. Interestingly, C5aR1^−/−^ mice had significantly higher levels of immune (CD45+) cells expressing IL-10 than WT mice even though the total percentage of immune infiltrate remained comparable between these two groups of mice **(Figure 3A and Supplementary Figure 2N)**. IL-10 signaling in CX3CR1+ macrophages has been recently found to be an important contributor to gut defense and homeostasis maintenance (Shouval et al., 2014; Zigmond et al., 2014). Therefore, we assessed the percentage of macrophages (F4/80+ and CX3CR1+) expressing IL-10 and found that C5aR1^−/−^ mice displayed increased levels of IL-10 expressing macrophages despite the total percentage of F4/80+ macrophages being comparable in C5aR1^−/−^ and WT mice **(Figures 3B, C and Supplementary 3E)**. We did note, however, a significant increase in F4/80+CX3CR1+ cells in the intestines of C5aR1^−/−^ mice but not in the spleen **(Supplementary Figure 3F-H).** We also, confirmed that, as expected, C5aR1^−/−^ mice did not express C5aR1 **(Supplementary Figure 3I).**

**Figure 3.**
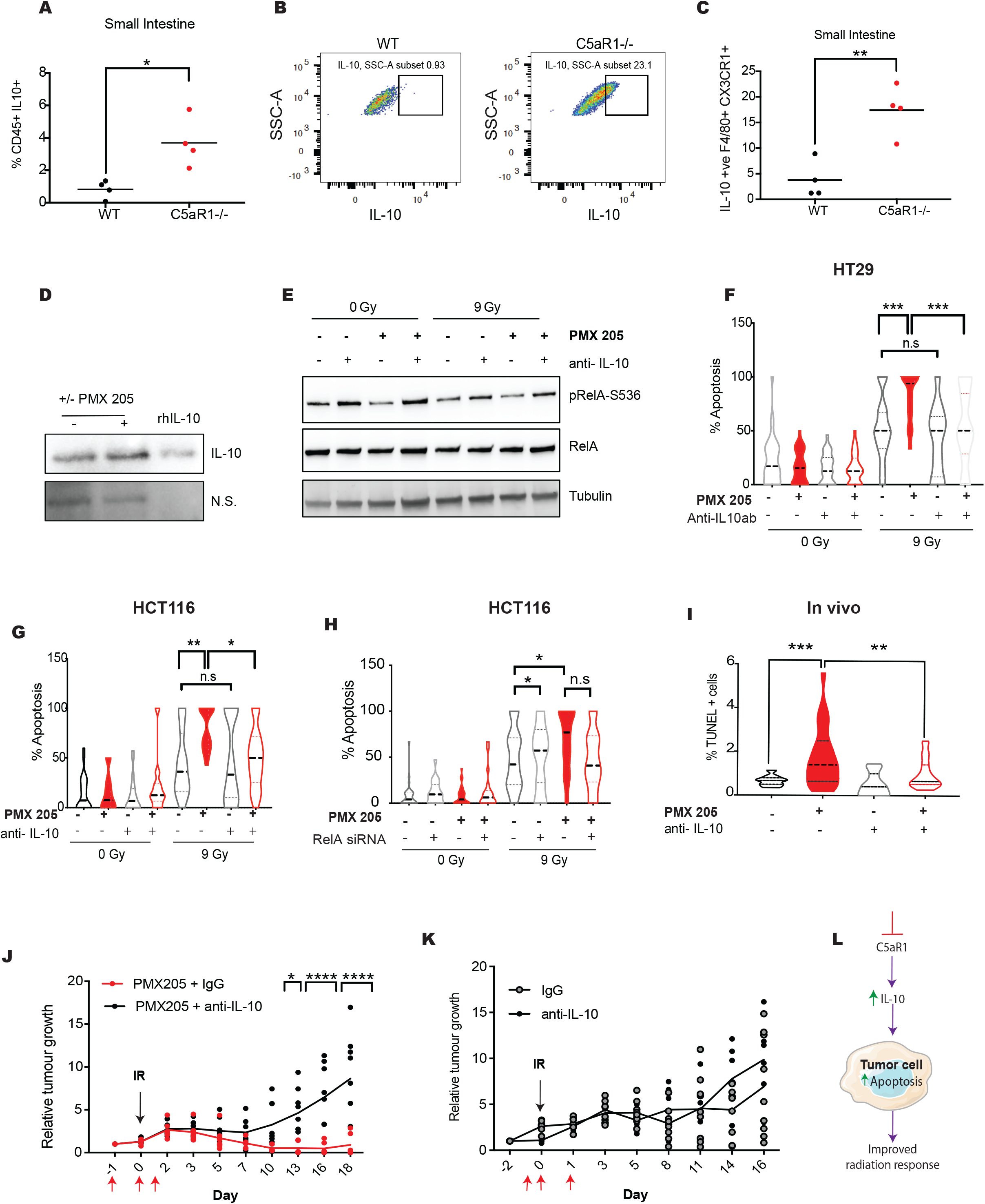
Targeting C5aR1 increases apoptosis and mediates improved tumor response in an IL-10-dependent manner. (A) Graph shows the % of CD45+ and IL-10+ cells found in small intestines of WT or C5aR1^−/−^ mice. * = p<0.05, 2-tailed t-test. Individual points represent individual mice per group. (B) Representative FACS plots showing the % IL-10 positivity in F4/80+ and CX3CR1+ cells found in small intestines one WT or C5aR1^−/−^ mouse respectively. (C) Graph shows the % IL-10 positivity in F4/80+ and CX3CR1+ cells found in small intestines of WT or C5aR1^−/−^ mice. ** = p<0.01, 2-tailed t-test. Individual points represent individual mice per group. (D) HCT 116 cells were treated with either vehicle or PMX205 in phenol and serum-free media. Western blotting was carried on condition media from these cells with the antibodies indicated. A non-specific band (denoted n.s) was used as an indication of loading. 80 ng of human recombinant IL-10 (rhIL-10) was used as a positive control. n=3. (E) HT29 cells were treated with either vehicle or PMX205 and either IgG or IL-10 blocking antibody from 1 hour before irradiation (IR) with either 0 or 9 Gy. Cells were harvested 48 hours post-IR. Western blotting was carried out with the antibodies indicated. Tubulin was used as the loading control. (F) The graph represents the number of apoptotic/non-apoptotic cells expressed as a % of the whole population for HT29 cells treated with either vehicle or PMX205 and either IgG or IL-10 blocking antibody from 1 hour before irradiation (IR) with either 0 or 9 Gy. Cells were harvested 48 hours post-IR. Independent fields of view are show, n=3. **** = p<0.0001 by 2-way ANOVA with Tukey-Kramer comparison test. (G) The graph represents the number of apoptotic/non-apoptotic cells expressed as a % of the whole population for HCT 116 cells treated with either vehicle or PMX205 and either IgG or IL-10 blocking antibody from 1 hour before irradiation (IR) with either 0 or 9 Gy. Cells were harvested 48 hours post-IR. n=3. **** = p<0.0001 by 2-way ANOVA with Dunnett’s comparison test. (H) The graph represents the number of apoptotic/non-apoptotic cells expressed as a % of the whole population for HCT 116 cells transfected with either Scr or RelA siRNA and treated with either vehicle or PMX205 from 1 hour before irradiation (IR) with either 0 or 9 Gy. Cells were harvested 48 hours post-IR. Independent fields of view from a representative experiment are show, n=3. ** = p<0.01, * = p<0.05 by 2-way ANOVA with Dunnett’s comparison test. (I) The graph represents the number of TUNEL positive (+) cells found in MC38 subcutaneous tumors treated with either vehicle or PMX205 and either IgG or IL-10 blocking antibody 1 day before irradiation (10 Gy), on the day of and 1 day post-irradiation. Individual points represent independent fields of view from up to 3 different tumors per group. **** = p<0.0001, n.s = not significant, 2-way ANOVA with Dunnett’s comparison test. (J) Relative tumor growth curves are shown for MC38 colorectal cancer cells treated with 10 Gy single dose irradiation, PMX205 treatment and either IgG or IL-10 blocking antibody for 3 doses flanking the irradiation dose (on day 0, 1 and 2). * = p<0.05, ** = p<0.01 **** = p<0.0001 by 2-way ANOVA. Individual points represent individual mice per group. (K) Relative tumor growth curves are shown for MC38 colorectal cancer cells treated with 10 Gy single dose irradiation, vehicle treatment and either IgG or IL-10 blocking antibody for 3 doses flanking the irradiation dose (on day 0, 1 and 2). Individual points represent individual mice per group. (L) Working hypothesis model: targeting C5aR1 increases both IL-10 expression in the small intestine and IL-10 secretion by tumor cells. IL-10 attenuates RelA phosphorylation and mediates increased tumor cell apoptosis following irradiation.

Having established the presence of increased IL-10 in the small intestines of C5aR1^−/−^ (compared to WT) mice and the increased expression of IL-10 in cell populations associated with intestinal injury defense, we next assessed levels of IL-10 secretion in colorectal cancer cells. We observed increased levels of IL-10 in conditioned media from colorectal cancer cells treated with PMX205 **(Figure 3D).** Increased IL-10 secretion in these cells was associated with increased STAT3 phosphorylation at tyrosine 705, suggesting active IL-10 signaling occurs in these cells (STAT3 signaling is thought to be activated downstream of IL-10 and we indeed observed reduced STAT3 phosphorylation following IL-10R knockdown)(Yu et al., 2007) **(Supplementary Figure 3J)**. Furthermore, in line with IL-10 having a role in modulating NF-κB signaling in cancer cells, we noted that blocking IL-10 abrogated the decrease in RelA phosphorylation observed following PMX205 treatment **(Figure 3E)**. Importantly, we confirmed that the effects of IL-10 on NF-κB signaling were tumor-dependent. In the intestines, we did not observe any significant differences in mRNA levels of downstream NF-κB target genes between WT and C5aR1^−/−^ mice, nor did we observe any IL-10 dependent changes in RelA phosphorylation (**Supplementary Figures 2L and 3L-N)**.

Next, we investigated the role of IL-10 on tumor radiation response. We hypothesized that the increase in apoptosis observed following PMX205 treatment was IL-10-dependent. To test this hypothesis, we treated colorectal cancer cells with PMX205 and IL-10 blocking antibody. As expected, treatment of colorectal cancer cells with PMX205 and irradiation resulted in increased apoptosis. In line with PMX205 inducing apoptosis in an IL-10-dependent manner, treatment with blocking IL-10 antibody together with PMX205 abrogated the increased apoptosis observed following C5aR1 inhibition **(Figure 3F and 3G)**. Consistent with our hypothesis that the apoptosis observed following PMX205 treatment is p53-independent, we observed the same effects in p53 WT cells (HCT 116) and p53 mutant cells (HT29) **(Figure 3F and G).** To confirm that the effects of PMX205 were occurring downstream of NF-κB signaling we depleted RelA and found that while RelA depletion increased apoptosis levels in vehicle treated cells following irradiation (as expected); there was no further increase in apoptosis in PMX205 treated cells (presumably since these cells already have reduced “active” RelA **(Figure 3H and Supplementary Figure 3O)**. Of note, the robust, IL-10-dependent apoptosis observed following C5aR1 inhibition was observed *in vitro*, reinforcing the notion that targeting C5aR1 can improve tumor radiation responses in the absence of a functional immune system (and therefore T-cell infiltration). Finally, we confirmed that the increased apoptosis observed *in vitro* resulted in similar effects in terms of apoptosis and decreased tumor growth *in vivo* in the same model where we had previously observed reduced CD8+ T cell infiltration. As previously observed, treatment of tumors with PMX205 resulted in a significant tumor growth delay and this was accompanied by increased apoptosis **(Figure 3I-K).** However, treatment of tumors with irradiation, IL-10 blocking antibody and PMX205 did not result in any significant change in tumor growth or apoptosis compared to IgG (or IL10 blocking antibody alone) treated tumors **(Figure 3I-K)**. These data suggest that the improved tumor response observed following PMX205 treatment is IL-10 dependent **(Figure 3L)**.

Together these data indicate that 1) targeting C5aR1 results in increased IL-10 levels in both healthy intestines and secreted by cancer cells, 2) IL-10 drives increased apoptosis and improved tumor radiation response following PMX205 treatment.

### Pharmacologically targeting C5aR1 improves normal tissue and tumor radiation response in a combined tumor and normal tissue murine model

Having established that targeting C5aR1 results in reduced signs of radiation-induced small bowel toxicity and improved radiation responses in separate tumor and normal tissue studies we decided to move to a combined tumor and normal tissue model. Few preclinical studies have attempted to assess tumor and normal tissue radiation responses in a single model, probably due to the technical difficulties associated with these experiments. We reasoned, however, that this approach would be essential for the future translation of any attempt to increase the therapeutic window of radiotherapy. For these studies we used a syngeneic orthotopic preclinical ovarian cancer model in which ID8 cells disseminate in the peritoneal cavity following injection (Roby et al., 2000). Total abdominal irradiation of tumor bearing mice then allows the assessment of both tumor and small intestine radiation responses. Importantly, increased C5aR1 expression is associated with reduced disease-free survival in ovarian cancer, suggesting that targeting C5aR1 might also be beneficial in this tumor type **(Figure 4A)**. Of note, the ID8 disseminated ovarian cancer model closely recapitulates the course of human disease in that disease progression is very quick and untreated mice quickly succumb to the disease. In order to assess disease progression, and identify a suitable timepoint for studying both tumor and normal tissue responses, we initially assessed stool weights as an indicator of general intestinal function (Taniguchi et al., 2014). We waited until mice developed signs of disease (as would occur in the clinical setting), before delivering 0 or 9 Gy irradiation with or without PMX205 treatment **(Figure 4B)**. As expected, due to the severity of the disease model, all mice had a steep decrease in stool weight by day five following display of symptoms. However, mice receiving radiation and PMX205 were able to increase their stool weights by day 9 **(Figure 4C)**. These data suggest that intestinal function is improved in PMX205 and irradiation treated mice due to either reduced tumor burden and/or intestinal regeneration. To investigate this possibility, we repeated the experiment and assessed tumor burden and signs of intestinal regeneration at day three after irradiation **(Figure 4D)**. Although this was an early timepoint for assessment of tumor radiation response, attempts at intestinal regeneration are frequently assessed around this timepoint (Metcalfe et al., 2014; Olcina and Giaccia, 2016; Sheng et al., 2020; Wei et al., 2016). We indeed observed that three days post irradiation the majority of mice showed signs of early regeneration **(Supplementary Figure 4)**. As a control, we ensured that all mice had the same level of crypt damage **(Figure 4E)**. Importantly, mice treated with 9 Gy and PMX205, had a higher proportion of areas involved in early regeneration with the majority of the intestine involved in regeneration compared to vehicle treated mice **(Figure 4F)**. Of note, the improved % regeneration score area was observed even when only sections displaying the same level of crypt damage and % area affected by crypt damage where compared in order to ensure that the changes in regeneration scores were not just reflecting changes in crypt damage area. Furthermore, PMX205 and 9 Gy treated mice displayed the best tumor responses compared to any other group **(Figure 4G and H)**. As we had observed in our separate tumor and normal tissue models, these data suggest that targeting C5aR1 can increase the therapeutic index of radiotherapy.

**Figure 4.**
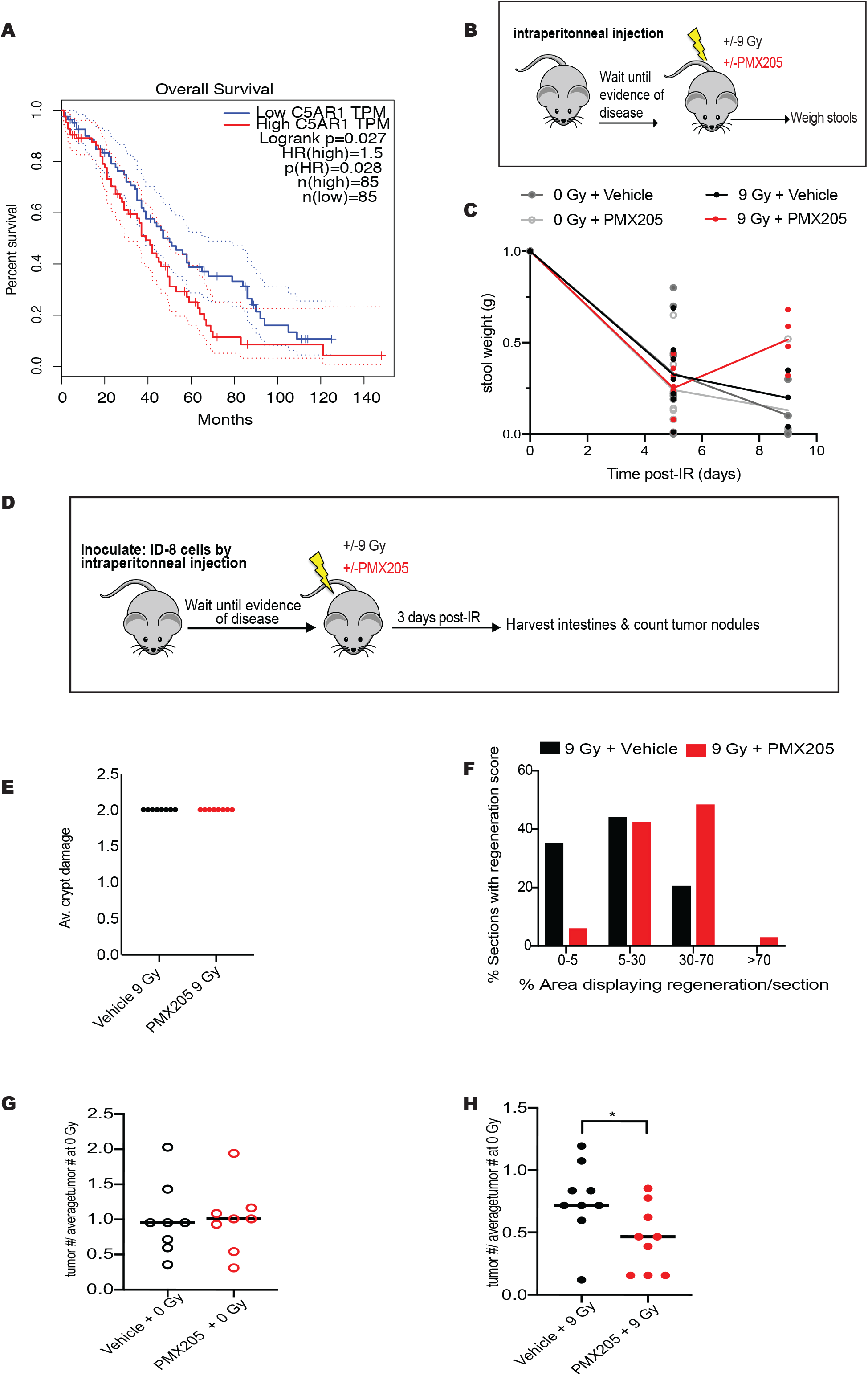
Pharmacologically targeting C5aR1 improves normal tissue and tumor radiation response in a combined tumor and normal tissue murine model. (A) GEPIA Kaplan-Meier (KM) curve for overall survival of TCGA ovarian cancer patients with high (red) or low (blue) C5AR1 mRNA expression levels is shown. Cutoff-High = 80%, Cutoff Low =20%. (http://gepia.cancer-pku.cn). (B) Schematic representation of the ovarian cancer model used to assess stool weights as a surrogate for intestinal function following irradiation (0 vs 9 Gy) and treatment with vehicle or PMX205. (C) Graph shows stool weight in grams (g) of mice treated with either 0 or 9 Gy and vehicle or PMX205. Individual points represent individual mice per group. (D) Schematic representation of the ovarian cancer model used to assess tumor and normal tissue responses following irradiation and treatment with vehicle or PMX205. (E) Graph shows average crypt damage scores for the small intestines of C57/BL6 mice treated with vehicle or PMX205 and irradiated with 9 Gy total abdominal irradiation. Intestines were harvested 3 days post irradiation. Individual points represent individual mice per group. (F) C57/BL6 mice bearing ID8 tumors were treated with vehicle or PMX205 and 9 Gy total abdominal irradiation. Graph shows % of areas involved in regeneration for intestinal sections showing early regeneration scores of 1. n=9 per group. (G) C57/BL6 mice bearing ID8 tumors were treated with vehicle or PMX205 and 0 Gy total abdominal irradiation. Graph shows tumor number/average tumor number at 0 Gy. Individual points represent individual mice per group. (H) C57/BL6 mice bearing ID8 tumors were treated with vehicle or PMX205 and 9 Gy total abdominal irradiation. Graph shows tumor number/average tumor number at 0 Gy. * = p<0.05, 2-tailed t-test. Individual points represent individual mice per group.

## Discussion

Here we report that targeting C5aR1 (pharmacologically or genetically) results in increased IL-10 levels in both colorectal cancer cells and in the mouse small intestine. We propose that increased IL-10 poises tumors to respond to the ionizing radiation insult by mediating an increased tumor-specific apoptotic response. Importantly, we show that C5ar1 antagonist PMX205 results in improved tumor radiation responses even in a context of reduced CD8+ T-cell infiltration, suggesting that PMX205 could be a tumor radiosensitizer in immune cold tumors with a paucity of T-cell infiltration. This is supported by our tumor studies in immunocompromised mice that showed significantly increased apoptosis following PMX205 treatment as well as by our *in vitro* data. We acknowledge, however, that we did not perform a functional assessment of immune cells in our tumor models. It is also important to note, that we do not rule out the contribution of the immune system to the overall anti-tumor effects of targeting C5aR1. Instead, we propose that increased IL-10-dependent apoptosis is a new mechanism that can contribute to the improved anti-tumor immune response modulated by targeting C5aR1; and that might be more prevalent at higher C5aR1 antagonist doses than the ones used in this study. Compelling data suggests that the C3aR/C5aR1/IL-10 signaling axis regulates T-cell mediated anti-tumor immunity, and indeed C3aR/C5aR1 have been proposed as a new class of immune checkpoint receptors (Ajona et al., 2017; Wang et al., 2016, 2019). Blocking C3aR/C5aR1 results in increased IL-10 production by anti-tumor effector T-cells leading to T-cell expansion and enhanced T-cell mediated anti-tumor responses (Wang et al., 2016). Our data indicate that increased IL-10 also has a positive effect on tumor response by regulating increased apoptosis following irradiation, even in the absence of T-cells (for example *in vitro* or in athymic nude mice). Future studies should be aimed at targeting C5aR1 to exploit both anti-tumor T-cell mediated cytotoxicity and increased tumor cell apoptosis for enhanced radiation responses.

The importance of modulating anti-tumor immune responses following C5aR1 targeting has been highlighted in previous studies and probably contributed to the reduced tumor burden observed here in C5aR1^−/−^ mice. The striking defect in tumor uptake displayed by C5aR1^−/−^ mice, also complicates the investigation of tumor radiation responses in this model and might have contributed to some of the previous conflicting reports on the effects of targeting the complement system on tumor radiation responses. Indeed, two studies have previously investigated the effects of radiation on local complement activation in the tumor and found increased expression of complement proteins following either fractionated or single dose irradiation (Elvington et al., 2014; Surace et al., 2015). These studies, however, reported seemingly contradictory results in terms of the effects of targeting complement on tumor radiation response. The first study showed that targeting complement with targeted inhibitor CR2-Crry (that blocks complement activation at the level of C3) together with localized fractionated radiotherapy increased apoptosis and inflammation in a lymphoma tumor model (Elvington et al., 2014). The improved tumor responses observed with CR2-Crry and irradiation were found to be neutrophil dependent. Furthermore, the authors reported increased dendritic cells (DC) and CD8+ T-cell infiltration in tumors treated with CR2-Crry and irradiation (Elvington et al., 2014). A later study investigating the effects of targeting complement receptors C3aR1 and C5aR1 together with single dose radiotherapy suggested that complement was critical for the anti-tumor effects of radiation (Surace et al., 2015). In this study the authors proposed that production of anaphylatoxins C3a and C5a and signaling through their receptors was required for dendritic cell maturation and CD8+ T-cell activation (Surace et al., 2015). It is important to note that both of these studies used different tumor models, fractionation schedules and pharmacological strategies for complement inhibition. The open controversy that still remains in the field underscores the importance of further studies directly comparing fractionation and single dose schedules and further exploring the potential for targeting complement in the context of radiotherapy. We have compared single dose and biologically equivalent fractionation doses in colorectal cancer models to assess the effect of targeting complement at the level of C5aR1. Our results show that targeting C5aR1 with PMX205 improves the radiation response of tumors in both the single dose and fractionation settings. As mentioned above, we focused on the effects of PMX205 on the tumor cell itself since we did not observe significant immune infiltration changes in our subcutaneous, single dose fraction tumor model. While we do not completely rule out the contribution of the immune system in our experiments, we note that targeting C5aR1 exerts a strong pro-apoptotic effect on tumor cells independent of the immune system. The contribution of these noncanonical functions may differ between models/following inhibition of the complement system at different levels which could also explain some of the discrepancies in the previous studies. Furthermore, the pharmacological approach used here, with short dosing with PMX205, allows formation of upstream complement activation products and signaling through C3aR1, likely maintaining higher levels of complement system activation than in some of the previous studies. Overall, this study, therefore, expands on our previous understanding of the effects of targeting complement and highlights the importance of investigating immune-independent roles for complement that can affect the response to cancer treatments. These data open up a new dimension in the study of complement beyond its traditional role as an innate immune pathway and into the area of autocrine cancer signaling within the context of radiotherapy.

Importantly, following total abdominal irradiation, C5aR1^−/−^ mice as well as mice treated with PMX205, display signs of reduced radiation-induced toxicity. We propose that the increased baseline levels of IL-10 observed following C5aR1 targeting “set-up” these mice to be protected from the detrimental small bowel toxicity effects associated with irradiation. The increased levels of CX3CR1 expressing macrophages in the small intestines of C5aR1^−/−^ mice might also contribute to the survival of these mice. IL-10 signaling in high CX3CR1-expressing macrophages polarizes them to an anti-inflammatory phenotype that could contribute to the reduced radiation-induced damage observed (Shouval et al., 2014; Zigmond et al., 2014). Furthermore, our data suggest that C5aR1 depletion results in opposite apoptosis effects in the small intestine and in tumors. Namely, apoptosis is increased following C5aR1 depletion in tumors while it decreases in intestinal crypts. The differences in apoptotic response downstream of C5aR1 could be explained by the seemingly opposite signaling changes observed in these two contexts. In tumor cells we observed decreased RelA phosphorylation. However, in the small intestines RelA phosphorylation did not decrease. Differences in the downstream effects of the C5aR1-IL-10 axis in tumors cells and intestinal epithelia might be explained due to the distinct apoptotic potential displayed by these two cellular populations. The intestinal epithelia undergoes constant regeneration and therefore has a strong physiologic and pathologic apoptotic potential (Delgado et al., 2016). In contrast, tumor cells evade apoptosis due to the activation of oncogenic signaling. These inherent differences may help explain why the same signaling axis results in divergent molecular signaling and ultimately the opposite downstream apoptotic response (Hanahan and Weinberg, 2011).

To test the utility of PMX205 in a setting allowing simultaneous assessment of tumor and normal tissue responses we made use of a syngeneic orthotopic preclinical ovarian cancer model (Roby et al., 2000). Ovarian cancer metastasises throughout the peritoneal cavity resulting in dismal prognostic outcomes for patients with advanced disease (Rai et al., 2014). Although abdominal irradiation was historically clinically considered for the treatment of disseminated ovarian cancer, concerns over small bowel toxicity limited its widespread use (despite the radiosensitivity of these tumors)(Rai et al., 2014). Chemotherapy is instead currently used following surgery, even though 70% of patients experience abdominal recurrence (Rai et al., 2014). Given the effectiveness of radiotherapy at controlling tumor response, finding therapeutic strategies that can limit radiation-induced toxicity, in the context of clinically-relevant ovarian cancer models, could result in an improved treatment modality for an area of unmet clinical need (Rai et al., 2014). Notably, we show that in a model of disseminated ovarian cancer, treatment with PMX205 and radiotherapy results in signs of reduced radiation-induced normal tissue toxicity without compromising tumor control.

Collectively, our results shown that the overall effect of targeting C5aR1 is improved tumor radiation response (without the need for concomitant increased CD8+ T-cell infiltration) and decreased radiation-induced small bowel toxicity providing a potential therapeutic strategy for increasing the therapeutic window for radiotherapy in different tumor contexts.

## Materials and Methods

### Cell lines and treatments

HCT 116 male adult human epithelial colorectal carcinoma cells originally purchased from ATCC^®^ were a kind gift from Ester Hammond (Oxford, UK). HT-29 female adult colorectal adenocarcinoma cells were purchased from ATCC^®^. ID8 cells were a kind gift from Erinn Rankin, (Stanford USA). Cells were grown in DMEM with 10% FBS, in a standard humidified incubator at 37°C and 5% CO_2_. CT26 (mouse colon carcinoma) cells were a kind gift from Matt Bogyo (Stanford, USA) and were grown in RPMI with 10% FBS, in a standard humidified incubator at 37°C and 5% CO_2_. CT26 is a N-nitroso-N-methylurethane (NNMU)-induced undifferentiated colon carcinoma cell line induced in BALB/c mice (Corbett et al., 1975; Wang et al., 1995). All cell lines were routinely tested for mycoplasma and found to be negative.

#### IL-10 blocking antibody treatment

Cells were pre-treated with 1 μg/ml IL-10 antibody (R&D # MAB217-100) 1 hour before irradiation. Mouse anti-human IgG control (Santa Cruz, #sc-2025) was used as the IgG control.

#### PMX205 treatment

For all *in vitro* experiments cells were treated with 10 μg/ml PMX205 (Tocris #5196) dissolved in 20% ethanol/water. Vehicle control in these experiments refers to 20% ethanol/water. Cells were pre-treated for 1 hour before irradiation.

### Animal studies

CT26 cells were injected intraperitoneally into either WT or C5aR1^−/−^ 6-8 week old BALBc/J female mice (JAXX). Two weeks later when WT mice started showing visible signs of disease (including bloating) mice were randomized into one of four groups (WT 0 Gy, C5aR1^−/−^ 0 Gy, WT 12 Gy, C5aR1^−/−^ 12 Gy). Tissues from BALBc/J injected with CT26 cells were harvested five days post irradiatiom. ID8 cells were injected intraperitoneally into 6-8 week old female C57/BL6 mice (JAXX). When mice started showing visible signs of disease (including bloating) mice were randomized into one of four groups (Vehicle 0 Gy, PMX205 0 Gy, Vehicle 9 Gy, PMX205 9 Gy). Mice were irradiated using the X-Rad SmART (Precision X-Ray Inc., North Branford, CT) as described below. MC38 cells (5 × 10^5^) were injected subcutaneously into 6-8 week old female C57/BL6 (JAXX) at a single dorsal site. All total abdominal irradiations were performed on anaesthetized animals (with the use of ketamine 100mg/kg/xylazine 20mg/kg) and using a 225 kVp cabinet X-ray system filtered with 0.5 mm Cu (at Comparative Medicine Unit, Stanford, CA). Growing tumors (average, 80-100 mm^3^) were treated with irradiation (either 9 Gy as a single dose or equivalent fractionated doses 3 × 4.45 Gy) and either vehicle or PMX205 for the days flanking the irradiation. In order to perform irradiations of subcutaneous tumors, mice were anesthetized in a knockdown chamber with a mixture of 3% isoflurane and 100% O_2_ and placed inside the irradiator cabinet on the subject stage. Anesthesia was maintained using 1.5% isoflurane in O_2_ delivered via a nose cone. Mice were monitored for breathing with a webcam fitted in the cabinet. The prescribed dose was calculated for a point chosen approximately in the middle of the tumor. The beam angle was chosen so as to target the superficial tumor while sparing critical organs within the mouse. The X-Rad SmART (Precision X-Ray Inc., North Branford, CT) was used for the irradiation of subcutaneous tumors. The X-ray tube and detector plate are mounted on a U shape gantry that rotates 360° in X and Y plane around the animal stage. The animal stage is supplied by a nose cone delivering isoflurane anesthesia and can move ±10cm in the X and Y directions and ±15cm in the Z direction. Pre-treatment computed tomography (CT) images were acquired to facilitate treatment planning, using a beam energy of 40kVp, a beam filter of 2 mm Al, and a voxel size of 0.2 or 0.1 mm. CT images collected were loaded onto RT_Image and a treatment plan was created using a single beam. The open-source RT_Image software package, version 3.13.1, running on IDL version 8.5.1 was used to visualize CT images and perform treatment planning (Graves et al., 2007). Therapeutic irradiations were performed using an x-ray energy of 225kVp, a current of 13mA, a power of 3000 Watts, and a beam filter of 0.3 mm Cu, producing a dose rate of ~300cGy/min at the isocenter. Treatment x-ray beams were shaped using a 10 or 15 mm collimator to selectively irradiate the target while sparing adjacent tissue when performing xenograft irradiation. For total abdominal irradiations square the 20 mm collimator with two beams; 0 and 180 degrees was used. After radiation delivery mice were removed and placed in a recovery box prior to being returned to their cage.

For experiments detailed in Figure 3, MC38 cells (5 × 10^5^) were injected subcutaneously into 6-8 week old female C57/BL6 (JAXX) at a single dorsal site. Growing tumors (average, 80-100 mm^3^) were treated with 10 Gy (as described above) and either PMX205 or vehicle and either IL-10 blocking antibody (LEAF Purified clone JES5-16E3, Biolegend #505012) (50 μg 24 hours before irradiation, 100 μg on the day of irradiation and 15 μg 24 hours post irradiation) or IgG control (LEAF Purified rat IgG2b, κ isotype control #400637, Biolegend) (same dosing and concentration as IL-10 antibody). WT or C5aR1^−/−^ mice were also treated with either IL-10 blocking antibody (Biolegend) (50 μg 24 hours before irradiation, 100 μg on the day or irradiation and 15 μg 24 hours post-irradiation) or IgG control (same dosing and concentration as IL-10 antibody).

For all experiments, tumors were measured with the use of calipers and volumes calculated by the ellipsoid estimation method as previously described (Taniguchi et al., 2014). For *in vivo* experiments involving PMX205 treatment, 10 mg/kg PMX205 (Tocris #5196, or synthesized and purified as previously described (Kumar et al., 2020)) was administered to mice orally flanking the irradiation doses. Vehicle control in these experiments refers to 20% ethanol/water.

Stools for each individual mouse (housed separately) were collected and weighed at the days indicated.

All procedures were performed in accordance with the National Institutes of Health (NIH) guidelines for the use and care of live animals (approved by the Stanford University Institutional Animal Care and Use Committee). Mice were euthanized as per APLAC guidelines in a CO_2_ chamber by cervical dislocation.

### Histology

Following euthanasia, small intestinal tissues were harvested and immersion-fixed in 10% neutral buffered formalin for 72 hours. Following fixation, serial longitudinal sections of small intestine were submitted for routine sectioning (Histo-Tec Laboratory, Inc. Hayward, CA). Briefly, slides were routinely processed, embedded in paraffin, section at 5 μm, and stained with hematoxylin and eosin (HE). To assess for radiation-induced toxicity, an ordinal grading scale was created that evaluated the following six parameters: 1) neutrophilic inflammation, 2) tissue area (%) affected by neutrophilic inflammation, 3) crypt damage, 4) tissue area (%) affected by crypt damage, 5) crypt regeneration impairment, and 6) tissue area (%) affected by crypt regeneration impairment (see Supplementary Table 2 for scoring definitions). Six sections of small intestine (duodenum, jejunum, and ileum) were evaluated per mouse. A total damage score was calculated for each mouse by adding individual scores across all six individually evaluated parameters (highest score possible = 18). All sections were evaluated by a single board-certified veterinary pathologist (K.M.C.).

### Immunohistochemistry

Formalin-fixed paraffin-embedded sections were dewaxed by soaking the slides in xylene for 15 minutes followed by 5 minute washes in 100%, 90%, 80%, and 70% ethanol, and 5 min in 3% hydrogen peroxide. The sections were then placed in citrate buffer (10mM citric acid, 0.05% Tween-20) and microwaved for 10 minutes for antigen retrieval. Quenching was performed with 1% hydrogen peroxide in methanol for 20 min. Sections were stained for Ki67 (Themo Fischer Scientific MA5-14520) or MECA32 (BD #BD553849). After incubation with secondary antibody— biotinylated anti-horse IgG (Vector Laboratories, BA-8000) and biotinylated anti-goat IgG (Vector Laboratories, BA-9500), avidin-biotin complex technique was performed with the ABC kit (Thermofisher, 32020) to enhance detection, which was accomplished using the DAB Substrate Kit (Abcam, ab64238) for 45 seconds. Counterstaining of nuclei was done with hematoxylin for 30 seconds then dehydrated with ethanol. Slides were mounted using DPX mounting medium and imaged. Stained slides were scanned with a NanoZoomer 2.0-RS Digital Slide Scanner (Hamamatsu). The two sections were overlaid and 250 μm x 250 μm squares were magnified individually. Image saturation was increased to 200% to increase contrast between the blue-stained nuclei and the brown-stained marker of interest. Staining was quantified by counting number of positive cells/crypts in 10 random high-power fields per slide.

#### TUNEL assay

ApopTag® (Millipore #S7100) was used to stain 4 mm formalin-fixed paraffin-embedded small intestinal or tumor sections. Stained slides were scanned with a NanoZoomer 2.0-RS Digital Slide Scanner (Hamamatsu). For tumor studies, fifteen randomly chosen fields of view at 10x magnification were selected for analysis via ImageJ software (NIH). Images were first converted to 8-bit black and white, then the threshold was adjusted to 0-200 to eliminate background signal. The remaining particles larger than 10 square pixels were counted. The % of counts/total area analyzed per field is shown. For small intestines, the number of TUNEL+ cells per crypt (or crypts and villi) were manually counted in at least 10 fields of view per section.

Apoptosis assessment by morphology in cell lines was carried out as previously described (Olcina et al., 2015; Olcina et al., 2020). Adherent and detached cells (in media) were collected and fixed in 4% PFA for 15 minutes at room temperature. PFA was then removed and cells were washed in 1 ml PBS. 10 μl of fixed cells in PBS was placed in each slide and mixed with ProLong™ Gold antifade mountant with DAPI (Invitrogen #P36935). A coverslip was placed on top of the sample and slides were left to dry overnight. Slides were imaged using a DSM6000, DMi8 or DMI6000 (Leica) microscope with 40x or 60x oil objectives. The number of cells with fragmented DNA and the total number of cells per field was counted (with typically at least 10 fields counted for every treatment).

### Immunoblotting

Cells were lysed in UTB (9 M urea, 75 mM Tris-HCl pH 7.5 and 0.15 M β-mercaptoethanol) and sonicated briefly before quantification as described in detail in (https://data.mendeley.com). Small intestinal tissue was harvested at the timepoints indicated in each corresponding figure legend and immediately placed in RNA*later*™ according to manufacturer’s instructions. Small intestines were lysed in RIPA lysis buffer and homogenized with the use of a douncer before quantification. Antibodies used were: β-actin (Sigma # A5441, concentration: 1:5000). hFAB™ Rhodamine Anti-Tubulin (Bio-Rad #12004166), RelA/p65 (Cell Signaling #3034, concentration 1:1000), pRelA/p65-S536 (Cell Signaling #3033T, concentration 1:1000), STAT3 (Cell Signaling #30835, concentration 1:1000), pSTAT3-Ty705 (Cell Signaling #9131, concentration 1:1000), p53-S15 (Cell Signaling, #9284 concentration 1:1000), total AKT (Upstate #14276, concentration 1:1000), AKT-S473 (Cell Signaling #9271S, concentration 1:1000) and anti-human IL-10 (R&D #MAB217-100, concentration 1:1000). The BioRad Chemidoc XRS system was used. In each case experiments were carried out in triplicate and a representative blot is shown unless otherwise stated.

#### Survival of mice after irradiation

Actuarial survival was calculated by the Kaplan-Meier method as previously described (Taniguchi et al., 2014).

#### Flow cytometry analyses

C5aR1^−/−^ and WT control mice or C57/BL6 MC38 tumor bearing mice were euthanized as per APLAC guidelines. The whole small intestine was washed in ice-cold PBS and the tissue was minced into 1 mm pieces. Tumor, spleen and small intestine tissue was digested into a single suspension using the murine tumor dissociation kit from Miltenyi Biotech (Auburn, CA) as per the manufacturer’s protocol. After RBC lysis, cells were re-suspended in PBS, counted and then stained with Zombie NIR (BioLegend, San Diego, CA) for live/dead cell discrimination. Nonspecific binding was blocked using an anti-mouse CD16/32 (BioLegend, San Diego, CA) antibody. Following which, cell surface staining was performed using fluorophore conjugated antimouse CD45.1 (30-F11), CD11b (M1/70), CD11c (N418), Ly6G (1A8), Ly6C (HK1.4), F4/80 (BM8), CX3CR1 (SA011F11), CD8 (53-6.7), CD4 (GK1.5) from Biolegend (San Diego, CA). After surface staining, cells were washed, fixed and permeabilized using BD Cytofix/Cytoperm kit and then stained for IL-10 expression (JES3-19F1, Biolegend). Flow cytometry was performed on LSR Fortessa (BD Biosciences) in the Radiation Oncology Dept. FACS facility. Cell acquisition was performed the following day with FACSDiva software on an LSR II flow cytometer (BD Biosciences, San Jose, CA) and analyzed with FlowJo software (Tree Star Inc., San Carlos, CA). Compensations were attained using Anti-Rat and anti-hamster compensation beads (BD Biosciences). For fixable live/dead staining, compensation was performed using ArC amine reactive compensation beads (BD Biosciences).

Gating schemes of dissociated tissues was as follows: Epithelial cells (ZNIR−CD45−), Immune cells-(ZNIR−CD45+), CD8 T-cells (ZNIR−CD45+CD8+), macrophages (ZNIR−CD45+CD11b+F4/80+), Neutrophils (ZNIR−CD45+CD11b+Ly6G+), Monocytes (ZNIR−CD45+CD11b+Ly6C+). For positive staining determination for IL-10 and CX3CR1, respective FMO controls were used.

#### ELISA

Following euthanasia (as per APLAC guidelines), small intestinal tissue from either WT or C5aR1^−/−^ mice was harvested, cleaned in ice-cold PBS (to remove intestinal contents) and immediately placed in Hank’s Balanced Salt Solution (HBSS) buffer on ice/4 ^o^C for 4 hours. Following centrifugation at 13000 g for 15 minutes at 4 ^o^C supernatants were placed in a clean tube. Murine IL-10 Mini TMB (PeproTech # 900-TM53) or ABTS (PeproTech # 900-M53) ELISA Development Kits were used according to manufacturer’s instructions.

### qRT-PCR

C5aR1^−/−^ and WT mice were euthanized as per APLAC guidelines. Small intestinal tissue was harvested, carefully but quickly cleaned (to remove intestinal contents) at the timepoints indicated in each corresponding figure legend and immediately placed in RNA*later*™ according to manufacturer’s instructions. RNA was extracted using Trizol (Invitrogen/Life Technologies,# 15596018). iScript cDNA synthesis kit (Bio-Rad, # 1708891) was used to reverse transcribe cDNA from total RNA according to manufacturer’s instructions. Relative mRNA levels were calculated using the standard curve methodology using a 7900HT Fast Real-Time PCR System. Primers used: 18S F: GTGGAGCGATTTGTCTGGTT, 18S R: ACGCTGAGCCAGTCAGTGTA, BIRC2 F: GCTGGCTTCTATTACATAGGGC, BIRC2 R: CTCAATCGAGCAGAGTGTGTC, XIAP F: CGAGCTGGGTTTCTTTATACCG, XIAP R: GCAATTTGGGGATATTCTCCTGT, BCL2L1 F: GACAAGGAGATGCAGGTATTGG, BCL2L1 R: TCCCGTAGAGATCCACAAAAGT.

### siRNA transfection

RelA (L-003533-00), C5aR1 (L-005442-00), IL10R (L-007925-00) or non-targeting RNAi negative control (Scramble, D-001810-10) (all from Dharmacon) were transfected into HCT 116 cells using Lipofectamine^**®**^RNAiMax transfection reagent (Invitrogen, #13778075) at a final concentration of 50 nM; according to the manufacturers’ instructions. Cells were harvested 72 hours post-transfection.

### *In silico* screen

Data was accessed through the Depmap portal (https://depmap.org from 10^th^ September 2020). Essentiality scores were calculated by dividing the number of dependent cell lines for each gene/total number of cell lines in either CRISPR-cas9 and RNAi screens. Genes described as complement system components, receptors, proteases and regulators (as reported in (Ricklin et al., 2010)) were queried. The calculated essentiality score was presented as a color in the heatmap. Green = No cell line is dependent on that gene and therefore the gene is not essential. Red = The majority of cell lines are dependent, or the gene is essential. Yellow/Orange = intermediate dependence or essentiality. The essentiality scale is shown at the bottom, left hand side of the heatmap. *ATR* was included in the screen a positive control. See Supplementary Table 1 for calculated raw values. Target tractability was assessed by accessing canSAR data as displayed on the Depmap portal. A gene was only considered a hit if it was “druggable” based on structural and ligand-based assessment. Genes that are hits and are druggable are shown in green. Red = not druggable based on structural or ligand-based assessment. C1R and CF1 were not included in the heatmap due to lack of CRISPR-Cas9 or RNAi data. C4A, C4B, VSIG4, C8A, C8B, CD93 and CR1 were not included due to their very low expression in the majority of tissues.

## Author contributions

Conceptualization, M.M.O. and A.J.G. Methodology M.M.O., S.M., D.K.M., R.K.K. and K.M.C. Writing original draft, M.M.O. Writing review and editing, M.M.O., A.J.G., D.K.N., and T.M.W. Investigation, M.M.O., S.M., D.K.N., and R.K.K. Formal analysis, M.M.O., D.K.N., R.K.K., K.M.C. and R.V.E. Resources, A.J.G., T.M.W. and M.S. Supervision M.M.O. and A.J.G. Funding acquisition, M.M.O., A.J.G. and M.S.

## Acknowledgements

This work was supported by NIH Grants CA-67166 and CA-197713, the Silicon Valley Foundation, the Sydney Frank Foundation and the Kimmelman Fund (AJG). MMO was a Cancer Research Institute Irvington Fellow supported by the Cancer Research Institute and Stiftung für Forschung, Medical Faculty, UZH. RKK was supported by Stanford ChEM-H Undergraduate Scholars Program. MS and MMO were supported by a project grant from the Swiss National Foundation (31003A_163141).

## Notes

### Competing Interest Statement

Prof Woodruff has previously consulted to Alsonex Pty Ltd, who are commercially developing PMX205. He holds no stocks, shares or other commercial interest in this company.

## References

Ajona, D., Ortiz-Espinosa, S., Moreno, H., Lozano, T., Pajares, M.J., Agorreta, J., Bértolo, C., Lasarte, J.J., Vicent, S., Hoehlig, K., et al. (2017). A combined PD-1/C5a blockade synergistically protects against lung cancer growth and metastasis. Cancer Discov. 7, 694–703.

Andreyev, H.J.N. (2007). Gastrointestinal Problems after Pelvic Radiotherapy: the Past, the Present and the Future. Clin. Oncol.

Ashburn, J.H., and Kalady, M.F. (2016). Radiation-Induced Problems in Colorectal Surgery. Clin. Colon Rectal Surg.

Bai, Y., Wang, W., Li, S., Zhan, J., Li, H., Zhao, M., Zhou, X.A., Li, S., Li, X., Huo, Y., et al. (2019). C1QBP Promotes Homologous Recombination by Stabilizing MRE11 and Controlling the Assembly and Activation of MRE11/RAD50/NBS1 Complex. Mol. Cell.

Baird, J.R., Friedman, D., Cottam, B., Dubensky, T.W., Kanne, D.B., Bambina, S., Bahjat, K., Crittenden, M.R., and Gough, M.J. (2016). Radiotherapy combined with novel STING-targeting oligonucleotides results in regression of established tumors. Cancer Res.

Baumann, M., Krause, M., Overgaard, J., Debus, J., Bentzen, S.M., Daartz, J., Richter, C., Zips, D., and Bortfeld, T. (2016). Radiation oncology in the era of precision medicine. Nat. Rev. Cancer 16, 234–249.

Bentzen, S.M. (2006). Preventing or reducing late side effects of radiation therapy: radiobiology meets molecular pathology. Nat. Rev. Cancer 6, 702–713.

Block, I., Müller, C., Sdogati, D., Pedersen, H., List, M., Jaskot, A.M., Syse, S.D., Lund Hansen, P., Schmidt, S., Christiansen, H., et al. (2019). CFP suppresses breast cancer cell growth by TES-mediated upregulation of the transcription factor DDIT3. Oncogene.

Burdette, D.L., Monroe, K.M., Sotelo-Troha, K., Iwig, J.S., Eckert, B., Hyodo, M., Hayakawa, Y., and Vance, R.E. (2011). STING is a direct innate immune sensor of cyclic di-GMP. Nature 478, 515–518.

Chen, C., Edelstein, L.C., and Gelinas, C. (2000). The Rel/NF-kappa B Family Directly Activates Expression of the Apoptosis Inhibitor Bcl-xL. Mol. Cell. Biol. 20, 2687–2695.

Cho, M.S., Vasquez, H.G., Rupaimoole, R., Pradeep, S., Wu, S., Zand, B., Han, H.D., Rodriguez-Aguayo, C., Bottsford-Miller, J., Huang, J., et al. (2014). Autocrine Effects of Tumor-Derived Complement. Cell Rep. 6, 1085–1095.

Cimprich, K.A., and Cortez, D. (2008). ATR: An essential regulator of genome integrity. Nat. Rev. Mol. Cell Biol.

Corbett, T.H., Griswold, D.P., Roberts, B.J., Peckham, J.C., and Schabel, F.M. (1975). Tumor Induction Relationships in Development of Transplantable Cancers of the Colon in Mice for Chemotherapy Assays, with a Note on Carcinogen Structure. Cancer Res.

Dar, T.B., Henson, R.M., and Shiao, S.L. (2019). Targeting innate immunity to enhance the efficacy of radiation therapy. Front. Immunol.

Delgado, M.E., Grabinger, T., and Brunner, T. (2016). Cell death at the intestinal epithelial front line. FEBS J.

Dewan, M.Z., Vanpouille-Box, C., Kawashima, N., DiNapoli, S., Babb, J.S., Formenti, S.C., Adams, S., and Demaria, S. (2012). Synergy of topical toll-like receptor 7 agonist with radiation and low-dose cyclophosphamide in a mouse model of cutaneous breast cancer. Clin. Cancer Res.

Duan, Q., Zhang, H., Zheng, J., and Zhang, L. (2020). Turning Cold into Hot: Firing up the Tumor Microenvironment. Trends in Cancer.

Egan, L.J., Eckmann, L., Greten, F.R., Chae, S., Li, Z., Myhre, G.M., Robine, S., Karin, M., and Kagnoff, M.F. (2004). I κB-kinaseβ-dependent NF-κ B activation provides radioprotection to the intestinal epithelium. Proc. Natl. Acad. Sci. U. S. A.

Elvington, M., Scheiber, M., Yang, X., Lyons, K., Jacqmin, D., Wadsworth, C., Marshall, D., Vanek, K., and Tomlinson, S. (2014). Complement-dependent modulation of antitumor immunity following radiation therapy. Cell Rep. 8, 818–830.

Flynn, R.L., and Zou, L. (2011). ATR: A master conductor of cellular responses to DNA replication stress. Trends Biochem. Sci.

Giaccia, A.J. (2014). Molecular Radiobiology: The State of the Art. J. Clin. Oncol. 32, 2871–2878.

Graves, E.E., Quon, A., and Loo, B.W. (2007). RT_image: An open-source tool for investigating PET in radiation oncology. Technol. Cancer Res. Treat.

Gros, P., Milder, F.J., and Janssen, B.J.C. (2008). Complement driven by conformational changes. Nat. Rev. Immunol. 8, 48–58.

Hanahan, D., and Weinberg, R.A. (2011). Hallmarks of cancer: The next generation. Cell 144, 646–674.

Hauer-Jensen, M., Denham, J.W., and Andreyev, H.J.N. (2014). Radiation enteropathy--pathogenesis, treatment and prevention. Nat. Rev. Gastroenterol. Hepatol. 11, 470–479.

Hofmeyer, T., Schmelz, S., Degiacomi, M.T., Dal Peraro, M., Daneschdar, M., Scrima, A., Van Den Heuvel, J., Heinz, D.W., and Kolmar, H. (2013). Arranged sevenfold: Structural insights into the C-terminal oligomerization domain of human C4b-binding protein. J. Mol. Biol.

Jain, U., Woodruff, T.M., and Stadnyk, A. (2013). The C5a receptor antagonist PMX205 ameliorates experimentally induced colitis associated with increased IL-4 and IL-10. Br. J. Pharmacol. 168, 488–501.

Jiang, X., Takahashi, N., Matsui, N., Tetsuka, T., and Okamoto, T. (2003). The NF-κB activation in lymphotoxin β receptor signaling depends on the phosphorylation of p65 at Serine 536. J. Biol. Chem.

Kavanagh, B.D., Pan, C.C., Dawson, L.A., Das, S.K., Li, X.A., Ten Haken, R.K., and Miften, M. (2010). Radiation Dose-Volume Effects in the Stomach and Small Bowel. Int. J. Radiat. Oncol. Biol. Phys.

Kumar, V., Lee, J.D., Clark, R.J., and Woodruff, T.M. (2018). Development and validation of a LC-MS/MS assay for pharmacokinetic studies of complement C5a receptor antagonists PMX53 and PMX205 in mice. Sci. Rep.

Kumar, V., Lee, J.D., Clark, R.J., Noakes, P.G., Taylor, S.M., and Woodruff, T.M. (2020). Preclinical Pharmacokinetics of Complement C5a Receptor Antagonists PMX53 and PMX205 in Mice. ACS Omega.

Lee, J.D., Kumar, V., Fung, J.N.T., Ruitenberg, M.J., Noakes, P.G., and Woodruff, T.M. (2017). Pharmacological inhibition of complement C5a-C5a1 receptor signalling ameliorates disease pathology in the hSOD1G93A mouse model of amyotrophic lateral sclerosis. Br. J. Pharmacol.

Leibowitz, B.J., Wei, L., Zhang, L., Ping, X., Epperly, M., Greenberger, J., Cheng, T., and Yu, J. (2014). Ionizing irradiation induces acute haematopoietic syndrome and gastrointestinal syndrome independently in mice. Nat. Commun.

Li, X.X., Lee, J.D., Massey, N.L., Guan, C., Robertson, A.A.B., Clark, R.J., and Woodruff, T.M. (2020). Pharmacological characterisation of small molecule C5aR1 inhibitors in human cells reveals biased activities for signalling and function. Biochem. Pharmacol.

Liang, H., Deng, L., Hou, Y., Meng, X., Huang, X., Rao, E., Zheng, W., Mauceri, H., Mack, M., Xu, M., et al. (2017). Host STING-dependent MDSC mobilization drives extrinsic radiation resistance. Nat. Commun.

Madrid, L. V., Mayo, M.W., Reuther, J.Y., and Baldwin, A.S. (2001). Akt Stimulates the Transactivation Potential of the RelA/p65 Subunit of NF-κB through Utilization of the IκB Kinase and Activation of the Mitogen-activated Protein Kinase p38. J. Biol. Chem.

Markiewski, M.M., DeAngelis, R.A., Benencia, F., Ricklin-Lichtsteiner, S.K., Koutoulaki, A., Gerard, C., Coukos, G., and Lambris, J.D. (2008). Modulation of the antitumor immune response by complement. Nat. Immunol. 9, 1225–1235.

Massard, C., Cassier, P., Bendell, J.C., Marie, D.B., Blery, M., Morehouse, C., Ascierto, M., Zerbib, R., Mitry, E., and Tolcher, A.W. (2019). Preliminary results of STELLAR-001, a dose escalation phase I study of the anti-C5aR, IPH5401, in combination with durvalumab in advanced solid tumours. Ann. Oncol.

Metcalfe, C., Kljavin, N.M., Ybarra, R., and de Sauvage, F.J. (2014). Lgr5+ Stem Cells Are Indispensable for Radiation-Induced Intestinal Regeneration. Cell Stem Cell 14, 149–159.

Moding, E.J., Kastan, M.B., and Kirsch, D.G. (2013). Strategies for optimizing the response of cancer and normal tissues to radiation. Nat. Rev. Drug Discov. 12, 526–542.

Olcina, M.M., and Giaccia, A.J. (2016). Reducing radiation-induced gastrointestinal toxicity -The role of the PHD/HIF axis. J. Clin. Invest. 126.

Olcina, M.M., Leszczynska, K.B., Senra, J.M., Isa, N.F., Harada, H., and Hammond, E.M. (2015). H3K9me3 facilitates hypoxia-induced p53-dependent apoptosis through repression of APAK. Oncogene.

Olcina, M.M., Kim, R.K., Balanis, N.G., Li, C.G., von Eyben, R., Graeber, T.G., Ricklin, D., Stucki, M., and Giaccia, A.J. (2020). Intracellular C4BPA Levels Regulate NF-κB-Dependent Apoptosis. IScience 23.

Olcina Monica, M., Kim Ryan, K., Balanis Nikolas, G., Li Caiyun, G., von Eyben, R., Graeber Thomas, G., Ricklin, D., Stucki, M., and Giaccia Amato, J. (2020). Intracellular C4BPA levels regulate NF-κB dependent apoptosis. IScience.

Pio, R., Ajona, D., and Lambris, J.D. (2013). Complement inhibition in cancer therapy. Semin. Immunol. 25, 54–64.

Rai, B., Bansal, A., Patel, F.D., and Sharma, S.C. (2014). Radiotherapy for ovarian cancers - redefining the role. Asian Pacific J. Cancer Prev.

Reis, E.S., Mastellos, D.C., Ricklin, D., Mantovani, A., and Lambris, J.D. (2017). Complement in cancer: untangling an intricate relationship. Nat. Rev. Immunol.

Ricklin, D., and Lambris, J.D. (2016). New milestones ahead in complement-targeted therapy. Semin. Immunol.

Ricklin, D., Hajishengallis, G., Yang, K., and Lambris, J.D. (2010). Complement: a key system for immune surveillance and homeostasis. Nat. Immunol. 11, 785–797.

Ricklin, D., Reis, E.S., and Lambris, J.D. (2016). Complement in disease: a defence system turning offensive. Nat. Rev. Nephrol.

Roby, K.F., Taylor, C.C., Sweetwood, J.P., Cheng, Y., Pace, J.L., Tawfik, O., Persons, D.L., Smith, P.G., and Terranova, P.F. (2000). Development of a syngeneic mouse model for events related to ovarian cancer. Carcinogenesis.

Roumenina, L.T., Daugan, M. V., Petitprez, F., Sautès-Fridman, C., and Fridman, W.H. (2019). Context-dependent roles of complement in cancer. Nat. Rev. Cancer.

Sayegh, E.T., Bloch, O., and Parsa, A.T. (2014). Complement anaphylatoxins as immune regulators in cancer. Cancer Med. 3, 747–758.

Sheng, X., Lin, Z., Lv, C., Shao, C., Bi, X., Deng, M., Xu, J., Guerrero-Juarez, C.F., Li, M., Wu, X., et al. (2020). Cycling Stem Cells Are Radioresistant and Regenerate the Intestine. Cell Rep.

Shouval, D.S., Biswas, A., Goettel, J.A., McCann, K., Conaway, E., Redhu, N.S., Mascanfroni, I.D., AlAdham, Z., Lavoie, S., Ibourk, M., et al. (2014). Interleukin-10 receptor signaling in innate immune cells regulates mucosal immune tolerance and anti-inflammatory macrophage function. Immunity.

Su, T., Zhang, Y., Valerie, K., Wang, X.Y., Lin, S., and Zhu, G. (2019). STING activation in cancer immunotherapy. Theranostics.

Surace, L., Lysenko, V., Fontana, A.O., Cecconi, V., Janssen, H., Bicvic, A., Okoniewski, M., Pruschy, M., Dummer, R., Neefjes, J., et al. (2015). Complement Is a Central Mediator of Radiotherapy-Induced Tumor-Specific Immunity and Clinical Response. Immunity 42, 767–777.

Suwinski, R., Wzietek, I., Tarnawski, R., Namysl-Kaletka, A., Kryj, M., Chmielarz, A., and Wydmanski, J. (2007). Moderately Low Alpha/Beta Ratio for Rectal Cancer May Best Explain the Outcome of Three Fractionation Schedules of Preoperative Radiotherapy. Int. J. Radiat. Oncol. • Biol. • Phys. 69, 793–799.

Taniguchi, C.M., Miao, Y.R., Diep, A.N., Wu, C., Rankin, E.B., Atwood, T.F., Xing, L., and Giaccia, A.J. (2014). PHD inhibition mitigates and protects against radiation-induced gastrointestinal toxicity via HIF2. Sci. Transl. Med. 6, 236ra64.

Voboril, R., and Weberova-Voborilova, J. (2007). Sensitization of colorectal cancer cells to irradiation by IL-4 and IL-10 is associated with inhibition of NF-kappaB. Neoplasma 54, 495–502.

Wang, M., Bronte, V., Chen, P.W., Gritz, L., Panicali, D., Rosenberg, S.A., and Restifo, N.P. (1995). Active immunotherapy of cancer with a nonreplicating recombinant fowlpox virus encoding a model tumor-associated antigen. J. Immunol.

Wang, Y., Meng, A., Lang, H., Brown, S.A., Konopa, J.L., Kindy, M.S., Schmiedt, R.A., Thompson, J.S., and Zhou, D. (2004). Activation of nuclear factor kappaB In vivo selectively protects the murine small intestine against ionizing radiation-induced damage. Cancer Res. 64, 6240–6246.

Wang, Y., Sun, S.N., Liu, Q., Yu, Y.Y., Guo, J., Wang, K., Xing, B.C., Zheng, Q.F., Campa, M.J., Patz, E.F., et al. (2016). Autocrine complement inhibits IL10-dependent T-cell-mediated antitumor immunity to promote tumor progression. Cancer Discov. 6, 1022–1035.

Wang, Y., Zhang, H., and He, Y.W. (2019). The Complement Receptors C3aR and C5aR Are a New Class of Immune Checkpoint Receptor in Cancer Immunotherapy. Front. Immunol.

Wei, L., Leibowitz, B.J., Xinwei, W., Epperly, M., Greenberger, J., Lin, Z., and Jian, Y. (2016). Inhibition of CDK4/6 protects against radiation-induced intestinal injury in mice. J. Clin. Invest.

Yu, H., Kortylewski, M., and Pardoll, D. (2007). Crosstalk between cancer and immune cells: Role of STAT3 in the tumour microenvironment. Nat. Rev. Immunol.

Zhong, H., May, M.J., Jimi, E., and Ghosh, S. (2002). The phosphorylation status of nuclear NF-κB determines its association with CBP/p300 or HDAC-1. Mol. Cell.

Zigmond, E., Bernshtein, B., Friedlander, G., Walker, C.R., Yona, S., Kim, K.W., Brenner, O., Krauthgamer, R., Varol, C., Müller, W., et al. (2014). Macrophage-restricted interleukin-10 receptor deficiency, but not IL-10 deficiency, causes severe spontaneous colitis. Immunity.

Zimmerer, T., Böcker, U., Wenz, F., and Singer, M. V. (2008). Medical prevention and treatment of acute and chronic radiation induced enteritis - Is there any proven therapy? A short review. Z. Gastroenterol.

