## Supplemental Files for "Targeting C5aR1 Increases the Therapeutic Window of Radiotherapy"

**Supplementary Table 1:** Raw data used in the generation of the heatmap shown in Figure 1A, see also methods section.

| Gene | CRISPR | RNAi | Druggable structure | Druggable by ligand based assessment |
|---|---|---|---|---|
| C2 | 0.001267427 | 0 | 0 | 1 |
| C3 | 0 | 0.001408451 | 0 | 0 |
| C5 | 0 | 0 | 0 | 0 |
| C6 | 0 | 0.004225352 | 0 | 0 |
| C7 | 0 | 0.021126761 | 0 | 0 |
| C8G | 0.012674271 | 0 | 0 | 1 |
| C9 | 0.001267427 | 0.001828154 | 0 | 0 |
| CR2 | 0.001267427 | 0 | 1 | 0 |
| C3AR1 | 0 | 0.007062147 | 1 | 0 |
| C5AR1 | 0 | 0 | 0 | 0 |
| C5AR2 | 0 | 0.004237288 | 1 | 0 |
| C1QBP | 0.60107095 | 0.01971831 | 1 | 1 |
| ITGAM | 0.001267427 | 0.015536723 | 0 | 0 |
| ITGAX | 0 | 0.005625879 | 0 | 1 |
| C4BPA | 0 | 0 | 0 | 0 |
| C4BPB | 0.001267427 | 0 | 1 | 1 |
| CPN1 | 0 | 0.003656307 | 1 | 1 |
| CPN2 | 0.007604563 | 0 | 1 | 0 |
| CALR | 0.015209125 | 0 | 0 | 1 |
| CLU | 0 | 0 | 1 | 1 |
| CFH | 0 | 0 | 0 | 1 |
| CD55 | 0.00390117 | 0.002816901 | 0 | 1 |
| CD59 | 0 | 0.049360146 | 1 | 0 |
| CD46 | 0.010403121 | 0.001408451 | 0 | 1 |
| VTN | 0.00887199 | 0.014577259 | 0 | 0 |
| SERPING1 | 0.001267427 | 0.004237288 | 1 | 1 |
| C1RL | 0 | 0 | 1 | 1 |
| C1S | 0 | 0.005649718 | 0 | 0 |
| MASP1 | 0 | 0.4092827 | 0 | 0 |
| MASP2 | 0.013941698 | 0.169491525 | 0 | 0 |
| CFB | 0.040557668 | 0 | 0 | 1 |
| CFD | 0.007604563 | 0 | 0 | 1 |
| ATR | 0.983523447 | 0.335674157 | 0 | 0 |

**Supplementary Table 2:** Description of the parameters used in the histologic scoring system described in Supplementary Figure 1.

Supplementary Table 1. Histopathologic Scoring Criteria for Radiation-Induced Small Intestinal Toxicity

| Parameter | Score 0 | Score 1 | Score 2 | Score 3 |
|---|---|---|---|---|
| Neutrophilic Inflammation | No inflammation | Small, focal, or widely-separated areas of inflammation limited to the lamina propria | Multifocal to coalescing areas of inflammation that extend into the submucosa | Transmural inflammatory infiltration |
| Tissue Area (%) Affected by Neutrophilic Inflammation | 0-5% involvement | 6-30% involvement | 31-70% involvement | >70% involvement |
| Crypt Damage | No crypt damage | Apoptotic crypt epithelial cells at any level of the crypt | Loss of proliferative crypt epithelial cells with residual Paneth cells | Complete crypt loss |
| Tissue Area (%) Affected by Crypt Damage | 0-5% involvement | 6-30% involvement | 31-70% involvement | >70% involvement |
| Crypt Regeneration Impairment | Complete regeneration or non-irradiated normal crypt epithelium | Atypical* crypt epithelial cells only | Flattened crypt epithelial cells with atypical* crypt epithelial cells | No attempt at tissue repair |
| Tissue Area (%) Affected by Crypt Regeneration Impairment | 0-5% involvement | 6-30% involvement | 31-70% involvement | >70% involvement |

* = karyomegaly, euchromatin, multiple nucleoli, increased numbers of mitotic figures

**Supplementary Figure 1: C5aR1 as a good therapeutic target**

**(A)** GEPIA Kaplan-Meier (KM) curve for overall survival of all combined TCGA cancers with either high (red) or low (blue) C5aR1 mRNA expression levels is shown. Group Cutoff = Median (http://gepia.cancer-pku.cn).

**(B)** Prognoscan KM curve for colorectal cancer patients with high (red) or low (blue) C5AR1 mRNA expression levels is shown. This analysis was based on the PrognoScan database (http://www.prognoscan.org/) using the publicly available Gene Expression Omnibus (http://www.ncbi.nlm.nih.gov/geo) with the accession numbers GSE 14333. Corrected p-value = 0.009386.

**(C)** Representative images of WT or C5aR1$^{-/-}$ mice intraperitoneally injected with CT26 colorectal cancer cells and treated with either 0 Gy or 12 Gy total abdominal irradiation. This dose of irradiation was chosen since it is expected to result in death in >90% of animals. White triangles indicate the location of large tumor nodules.

**(D)** Graph shows the tumor weight (g) of CT26 cells injected intraperitoneally into WT or C5aR1$^{-/-}$ BALBc/J mice. Individual dots represent separate mice per group.

**(E)** Graph shows the tumor weight (g) of CT26 cells injected intraperitoneally into WT or C5aR1$^{-/-}$ BALBc/J mice irradiated with 12 Gy. Individual dots represent separate mice per group. * = p<0.01, 2 tailed t-test.

**(F)** Graph shows the weight of mice treated with 9 Gy single dose irradiation and either vehicle or PMX205 treatment for 3 doses flanking the irradiation dose (on day 0, 1 and 2). Individual lines represent individual mice per group

**(G)** Graph shows an increase in the total histologic damage score with increasing irradiation (IR) doses. Total histologic damage score obtained by adding scores of 6 individual histologic parameters. Intestines were harvested 2 days post-IR. See also Supplementary Table 1. **** = p<0.0001, *** = p<0.001 by 1 way ANOVA with Dunnett's comparison test. Individual points represent individual mice per group.

**(H)** Graph shows total histologic damage score of WT and C5aR1$^{-/-}$ mice. Intestines were harvested 2 days post irradiation. **** = p<0.0001, 2 tailed t-test.

**(I)** Graph shows the crypt damage scores for the duodenum of WT and C5aR1$^{-/-}$ mice receiving 9 Gy total abdominal irradiation (IR). Intestines were harvested 2 days post-IR. Individual points represent individual mice per group.

**(J)** Graph shows the crypt damage scores for the jejunum of WT and C5aR1$^{-/-}$ mice receiving 9 Gy total abdominal irradiation (IR). Intestines were harvested 2 days post-IR. Individual points represent individual mice per group.

**(K)** Graph shows the crypt damage scores for the ileum of WT and C5aR1$^{-/-}$ mice receiving 9 Gy total abdominal irradiation (IR). Intestines were harvested 2 days post-IR. Individual points represent individual mice per group.

**(L)** Graph shows the number of vessels/panel in WT or C5aR1$^{-/-}$ mice receiving 9 Gy total abdominal irradiation (IR). Intestines were harvested 3 days post-IR. Individual points represent individual mice per group.

**(M)** Representative image of KI67 immunoreactivity in a C5aR1$^{-/-}$ section is shown. Ki67-positive cells are those with dark brown nuclear immunoreactivity.

**(N)** Graph shows the % of Ki67+ cells in crypts of WT or C5aR1$^{-/-}$ mice receiving 9 Gy total abdominal irradiation (IR). Intestines were harvested 3 days post-IR. Individual points represent individual mice per group.

**(O)** Graph shows the number of TUNEL+ cells in the villi of WT or C5aR1$^{-/-}$ mice receiving 9 Gy total abdominal irradiation (IR). Intestines were harvested 2 days post-IR. Individual points represent individual mice per group.

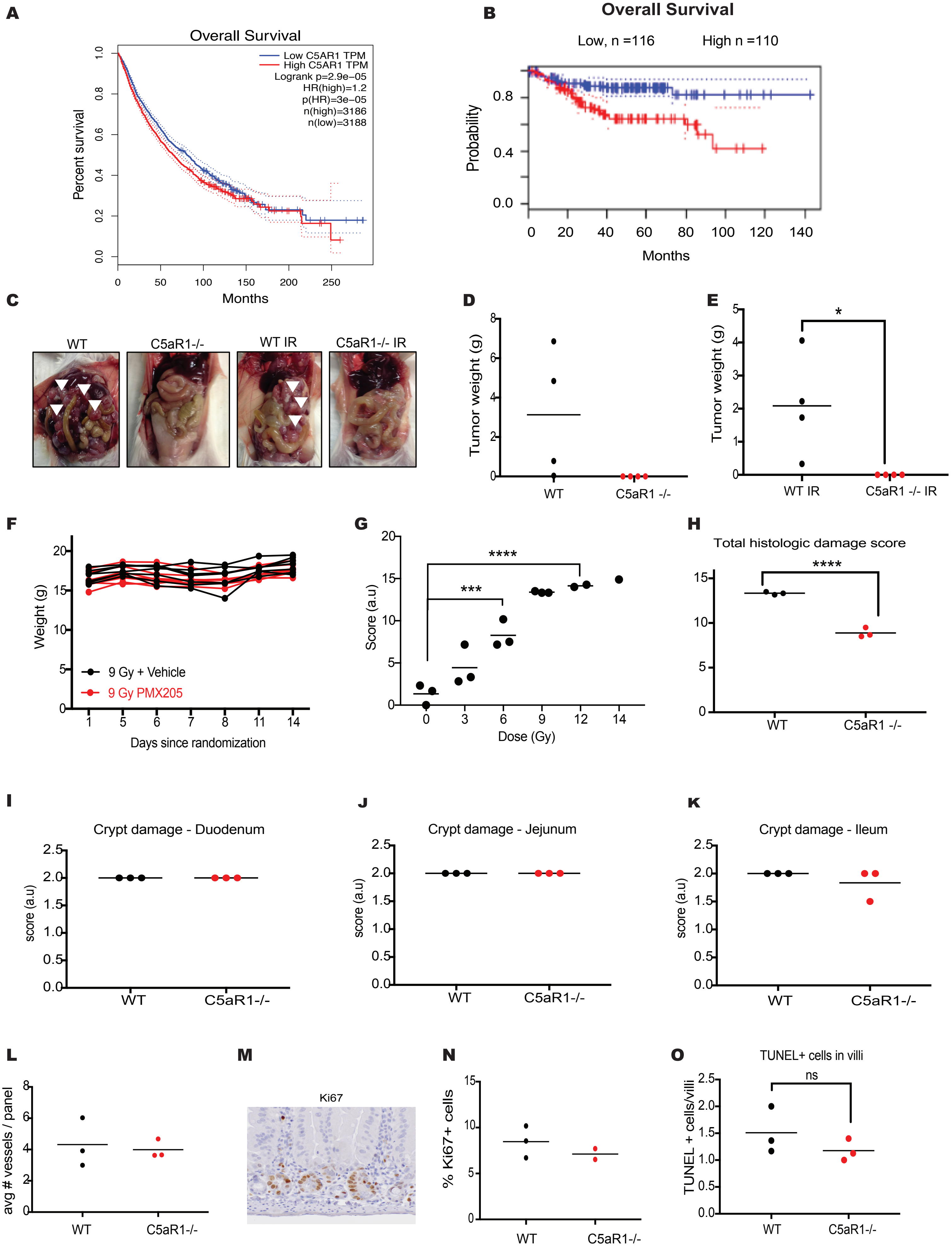

A
Overall Survival
Low C5AR1 TPM
High C5AR1 TPM
Logrank p=2.9e−05
HR(high)=1.2
p(HR)=3e−05
n(high)=3186
n(low)=3188
Percent survival
Months
B
Overall Survival
Low, n =116
High n =110
Probability
Months
C
WT
C5aR1-/-
WT IR
C5aR1-/- IR
D
Tumor weight (g)
WT
C5aR1 -/-
E
Tumor weight (g)
WT IR
C5aR1 -/- IR
F
Weight (g)
9 Gy + Vehicle
9 Gy PMX205
Days since randomization
G
Score (a.u)
Dose (Gy)
H
Total histologic damage score
WT
C5aR1 -/-
I
Crypt damage - Duodenum
score (a.u)
WT
C5aR1-/-
J
Crypt damage - Jejunum
score (a.u)
WT
C5aR1-/-
K
Crypt damage - Ileum
score (a.u)
WT
C5aR1-/-
L
avg # vessels / panel
WT
C5aR1-/-
M
Ki67
N
% Ki67+ cells
WT
C5aR1-/-
O
TUNEL+ cells in villi
ns
TUNEL + cells/villi
WT
C5aR1-/-

**Supplementary Figure 2. Targeting C5aR1 can mediate an improved radiation response in the context of reduced CD8+ T-cell infiltration**

- **(A)** Gating strategy for the different immune populations observed in the tumor
- **(B)** Graph shows CD3+ T-cells (as a % of live cells) in spleens of mice receiving 0 or 9 Gy and either vehicle or PMX205 treatment following the same dosing scheme as shown in Figure 1F. Tumors were harvested 7 days after irradiation with either 0 or 9 Gy. Individual points represent individual mice per group.
- **(C)** Graph shows NK cells (as a % of live cells) in spleens of mice receiving 0 or 9 Gy and either vehicle or PMX205 treatment following the same dosing scheme as shown in Figure 1F. Tumors were harvested 7 days after irradiation with either 0 or 9 Gy. Individual points represent individual mice per group.
- **(D)** Graph shows CD11b+ cells (as a % of live cells) in spleen of mice receiving 0 or 9 Gy and either vehicle or PMX205 treatment following the same dosing scheme as shown in Figure 1F. Tumors were harvested 7 days after irradiation with either 0 or 9 Gy. Individual points represent individual mice per group.
- **(E)** Graph shows M-MDSC cells (as a % of live cells) in spleen of mice receiving 0 or 9 Gy and either vehicle or PMX205 treatment following the same dosing scheme as shown in Figure 1F. Tumors were harvested 7 days after irradiation with either 0 or 9 Gy. Individual points represent individual mice per group.
- **(F)** Graph shows PMN-MDSC cells (as a % of live cells) in spleens of mice receiving 0 or 9 Gy and either vehicle or PMX205 treatment following the same dosing scheme as shown in Figure 1F. Tumors were harvested 7 days after irradiation with either 0 or 9 Gy. Individual points represent individual mice per group.
- **(G)** Relative tumor growth curves are shown for HCT 116 xenograft tumors grown in athymic nude mice treated with 9 Gy single dose irradiation and either vehicle or PMX205 treatment for 3 doses flanking the irradiation dose (on day 0, 1 and 2). **** = $p<0.0001$ by mixed effect analysis and comparing vehicle and PMX205 treated mice at day 33 and 35 by multiple by Sidak's multiple comparisons test.
- **(H)** HCT 116 cells were treated with 0 or 9 Gy and either vehicle or PMX205 from 1 hour before irradiation (IR). Cells were harvested 24 hours post-IR. Western blotting was carried with the antibodies indicated. β-actin was used as the loading control.
- **(I)** HCT 116 cells treated with either Scr or C5aR1 siRNA. Western blotting was carried with the antibodies indicated. β-actin was used as the loading control. n=2.
- **(J)** Protein levels of pRelA/total RelA in intestines of WT or C5aR1$^{-/-}$ mice treated with IgG and 9 Gy total abdominal irradiation (IR) is shown. Intestines were harvested 1 hour post-IR. Individual points represent individual mice per group.

**(K)** Protein levels of pRelA /β-actin in intestines of WT or C5aR1$^{-/-}$ mice treated with either IgG and 9 Gy total abdominal irradiation (IR) is shown. Intestines were harvested 1 hour post-IR.

**(L)** Western blot quantified in J and K is shown. WT or C5aR1$^{-/-}$ mice were treated with either IgG or IL-10 blocking antibody 1 day before irradiation (9 Gy), on the day of and 1 day post-irradiation. Small intestines were harvested 1 hour after irradiation. Individual lanes represent different mice per group. Western blotting was carried out with the antibodies indicated. β-actin was used as the loading control.

**(M)** Graph shows the % of CD45+ cells found in small intestine of WT or C5aR1$^{-/-}$ mice. Individual points represent individual mice per group.

**(N)** Graph shows CD8+ T-cells (as a % of live cells) in small intestine of WT or C5aR1$^{-/-}$ mice. Individual points represent individual mice per group. * = $p<0.01$, 2 tailed t-test.

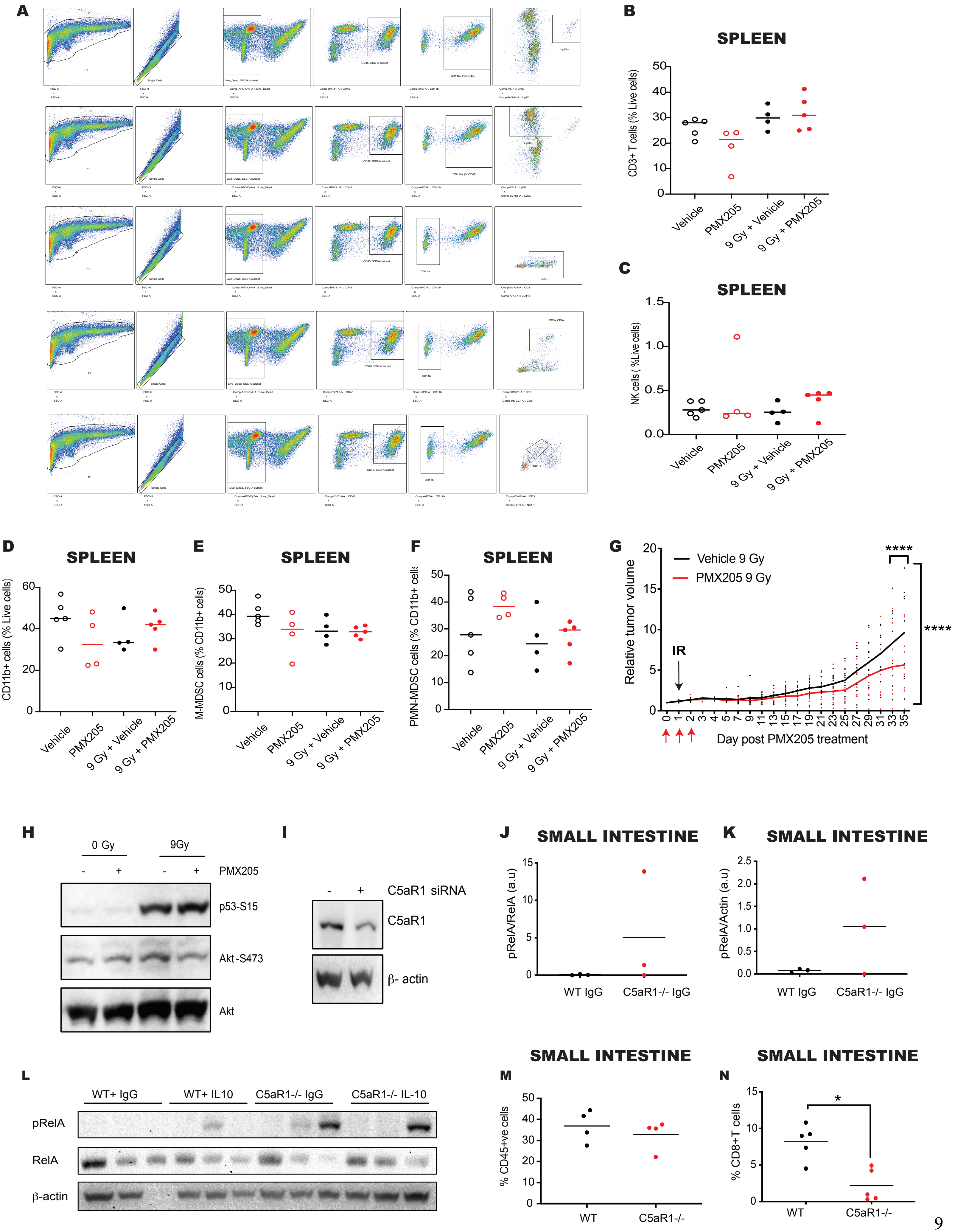
A
B
SPLEEN
CD3+ T cells (% Live cells)
Vehicle
PMX205
9 Gy + Vehicle
9 Gy + PMX205
C
SPLEEN
NK cells ( %Live cells)
D
SPLEEN
CD11b+ cells (% Live cells)
E
SPLEEN
M-MDSC cells (% CD11b+ cells)
F
SPLEEN
PMN-MDSC cells (% CD11b+ cells
G
Vehicle 9 Gy
PMX205 9 Gy
****
****
IR
Relative tumor volume
Day post PMX205 treatment
H
0 Gy
9Gy
PMX205
p53-S15
Akt -S473
Akt
I
C5aR1 siRNA
C5aR1
β- actin
J
SMALL INTESTINE
pRelA/RelA (a.u)
WT IgG
C5aR1-/- IgG
K
SMALL INTESTINE
pRelA/Actin (a.u)
WT IgG
C5aR1-/- IgG
L
WT+ IgG
WT+ IL10
C5aR1-/- IgG
C5aR1-/- IL-10
pRelA
RelA
β-actin
M
SMALL INTESTINE
% CD45+ve cells
WT
C5aR1-/-
N
SMALL INTESTINE
% CD8+T cells
*
WT
C5aR1-/-

**Supplementary Figure 3. Targeting C5aR1 increases apoptosis and mediates improved tumor response in an IL-10-dependent manner**

- **(A)** HCT 116 cells were treated with 0 or 9 Gy and either vehicle or PMX205 from 1 hour before irradiation (IR). Cells were harvested 48 hours post-IR. Western blotting was carried with the antibodies indicated. Tubulin was used as the loading control.
- **(B)** Graph shows the concentration of secreted IL-10 found in small intestine of WT or C5aR1$^{-/-}$ mice as measured by 2 different ELISA methods. Circles denote samples analyzed by ABTS and triangles denote samples analyzed by TMB method.
- **(C)** Graph shows the concentration of secreted IL-10 found in small intestines of WT or C5aR1$^{-/-}$ mice following 9 Gy total abdominal irradiation as measured by 2 different ELISA methods. Intestines were harvested 72 hours post-IR. Circles denote samples analyzed by ABTS and triangles denote samples analyzed by TMB method. Each sample represents an individual mouse.
- **(D)** Graph shows the % of CD45- IL10+ cells found in small intestine of WT or C5aR1$^{-/-}$ mice. Individual points represent individual mice per group.
- **(E)** Graph shows the % of F4/80+ cells found in small intestine of WT or C5aR1$^{-/-}$ mice. Individual points represent individual mice per group.
- **(F)** Graph shows the % of CX3CR1+ macrophages found in small intestines of WT or C5aR1$^{-/-}$ mice. Individual points represent individual mice per group.
- **(G)** Graph shows the % of CX3CR1+ F4/80 macrophages found in spleens of WT or C5aR1$^{-/-}$ mice. Individual points represent individual mice per group.
- **(H)** Gating strategy for F4/80+ CX3CR1- population is shown.
- **(I)** Graph shows the % C5aR1 positivity found in spleen of WT or C5aR1$^{-/-}$ mice. Individual points represent individual mice per group.
- **(J)** HCT 116 cells transfected with either Scr or IL-10R siRNA were treated with either vehicle or PMX205 from 1 hour before irradiation (IR). Cells were harvested 24 hours post-IR. Western blotting was carried with the antibodies indicated. $\beta$-actin was used as the loading control.
- **(K)** mRNA expression of *BIRC2/18S* in intestines of WT or C5aR1$^{-/-}$ mice treated with either IgG or IL-10 blocking antibody and 9 Gy total abdominal irradiation (IR) is shown. Intestines were harvested 72 hours post-IR. Individual points represent individual mice per group.
- **(L)** mRNA expression of *XIAP/18S* in intestines of WT or C5aR1$^{-/-}$ mice treated with either IgG or IL-10 blocking antibody and 9 Gy total abdominal irradiation (IR) is shown. Intestines were harvested 72 hours post-IR. Individual points represent individual mice per group.
- **(M)** mRNA expression of *BCL2L1/18S* in intestines of WT or C5aR1$^{-/-}$ mice treated with either IgG or IL-10 blocking antibody and 9 Gy total abdominal irradiation (IR) is shown. Intestines were harvested 72 hours post-IR. Individual points represent individual mice per group.

**(N)** HCT 116 cells were transfected with either Scr or RelA siRNA. Western blotting was carried with the antibodies indicated. β-actin was used as the loading control.

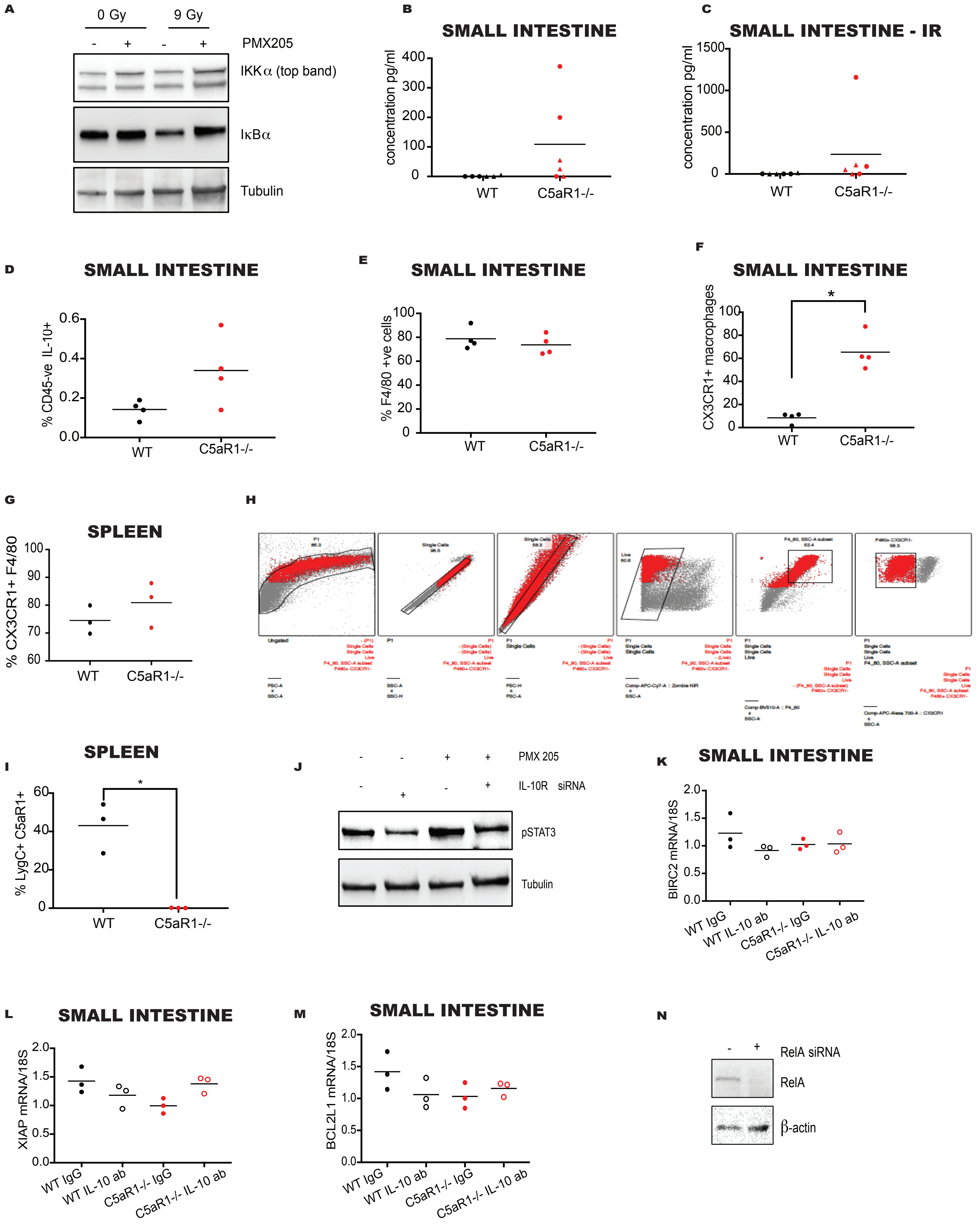
A
0 Gy
9 Gy
- + - +
PMX205
IKKα (top band)
IκBα
Tubulin
B
SMALL INTESTINE
concentration pg/ml
400
300
200
100
0
WT
C5aR1-/-
C
SMALL INTESTINE - IR
concentration pg/ml
1500
1000
500
0
WT
C5aR1-/-
D
SMALL INTESTINE
% CD45-ve IL-10+
0.6
0.4
0.2
0.0
WT
C5aR1-/-
E
SMALL INTESTINE
% F4/80 +ve cells
100
80
60
40
20
0
WT
C5aR1-/-
F
SMALL INTESTINE
CX3CR1+ macrophages
*
100
80
60
40
20
0
WT
C5aR1-/-
G
SPLEEN
% CX3CR1+ F4/80
100
90
80
70
60
WT
C5aR1-/-
H
I
SPLEEN
% LygC+ C5aR1+
*
60
40
20
0
WT
C5aR1-/-
J
- - + +
PMX 205
- + - +
IL-10R siRNA
pSTAT3
Tubulin
K
SMALL INTESTINE
BIRC2 mRNA/18S
2.0
1.5
1.0
0.5
0.0
WT IgG
WT IL-10 ab
C5aR1-/- IgG
C5aR1-/- IL-10 ab
L
SMALL INTESTINE
XIAP mRNA/18S
2.0
1.5
1.0
0.5
0.0
WT IgG
WT IL-10 ab
C5aR1-/- IgG
C5aR1-/- IL-10 ab
M
SMALL INTESTINE
BCL2L1 mRNA/18S
2.0
1.5
1.0
0.5
0.0
WT IgG
WT IL-10 ab
C5aR1-/- IgG
C5aR1-/- IL-10 ab
N
- +
RelA siRNA
RelA
β-actin

**Supplementary Figure 4. Pharmacologically targeting C5aR1 improves normal tissue and tumor radiation response in a combined tumor and normal tissue murine model**

C57/BL6 mice bearing ID8 tumors were treated vehicle or PMX205 and 9 Gy total abdominal irradiation. Graph shows early regeneration scores for mice treated with 9 Gy and either vehicle or PMX205. Regeneration score of 1 = Atypical cuboidal epithelial cells defined as cells presenting signs of attempted regeneration/repair such as karyomegaly, visible euchromatin, multiple nucleoli or increased numbers of mitotic figures. n = 9 mice/group.

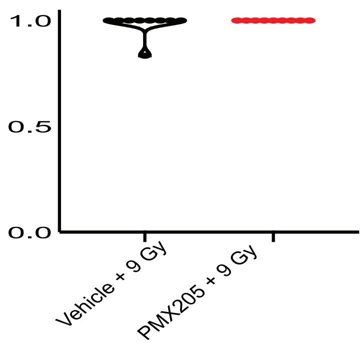
1.0
0.5
0.0
Vehicle + 9 Gy
PMX205 + 9 Gy
